# Stuck on a small tropical island: wide *in-situ* diversification of an urban-dwelling bat

**DOI:** 10.1101/2023.06.22.546033

**Authors:** Samantha Aguillon, Clara Castex, Avril Duchet, Magali Turpin, Gildas Le Minter, Camille Lebarbenchon, Axel O. G. Hoarau, Céline Toty, Léa Joffrin, Pablo Tortosa, Patrick Mavingui, Steven M. Goodman, Muriel Dietrich

## Abstract

Bats are often the only mammals naturally colonizing isolated islands and are thus an excellent model to study evolutionary processes of insular ecosystems. Here, we studied the Reunion free-tailed bat (*Mormopterus francoismoutoui*), an endemic species to Reunion Island that has adapted to urban settings. At regional scale, we investigated the evolutionary history of *Mormopterus* species, as well as on Reunion Island sex-specific and seasonal patterns of genetic structure. We used an extensive spatio-temporal sampling including 1,136 individuals from 18 roosts and three biological seasons (non-reproductive/winter, pregnancy/summer, and mating), with additional samples from *Mormopterus* species from neighbouring islands (*M. jugularis* of Madagascar and *M. acetabulosus* of Mauritius). Complementary information gathered from both microsatellite and mitochondrial markers revealed a high genetic diversity but no signal of spatial genetic structure and weak evidence of female philopatry. Regional analysis suggests a single colonization event for *M. francoismoutoui*, dated around 175,000 years ago, and followed by *in-situ* diversification and the evolution of divergent ancestral lineages, which today form a large metapopulation. Population expansion was relatively ancient (55,000 years ago) and thus not linked to human colonization of the island and the availability of new anthropic day-roost sites. Discordant structure between mitochondrial and microsatellite markers suggests the presence of yet-unknown mating sites, or the recent evolution of putative ecological adaptations. Our study illustrates how understanding mechanisms involved in speciation can be challenging and the importance of both mitochondrial and nuclear DNA in resolving the wide *in-situ* diversification of an urban-dwelling bat, endemic to a small island.

## Introduction

Islands have long been recognized as natural laboratories for studying evolution and biogeography. Their geographic isolation can lead to the evolution of unique adaptations in wildlife, with high levels of speciation and endemism (Warren et al., 2015). Island endemic bats represent a fascinating group of mammals that have colonized islands across numerous areas of the world. Given their ability to fly long distances, bats are often the only mammals naturally colonizing islands and are thus an excellent model group to examine evolutionary processes that can occur in insular ecosystems (Jones et al., 2009).

Once established on islands, bats face a diversity of selective pressures that can influence their evolution and diversification. Indeed, due to reduced species richness on islands with small surface areas, such insular ecosystems may lack predators or competitors and offer open ecological niches, which can favour the expansion of bat populations (Salinas-Ramos et al., 2020). On the contrary, insular ecosystems often have limited resources and can be subject to extreme events, such as volcanic activity, hurricanes or droughts, which can reduce bat populations (Calderón-Acevedo et al., 2021; Jones et al., 2001). Further, islands are vulnerable ecosystems that are highly susceptible to recent human-associated global changes, such as sea level rise and invasion by non-native species (Bellard et al., 2014). Due to their geographic isolation and limited dispersal opportunities, island endemic bat species may be particularly exposed to adverse effect of climate change (Festa et al., 2023). For example, a recent study on a Mediterranean island, the endemic Sardinian long-eared bat (*Plecotus sardus*, family Vespertilionidae) revealed a dramatic crash in population size, potentially due to recurrent wildfires and extreme temperatures (Ancillotto et al., 2021). Also, recent urbanization of island ecosystems could negatively affect the ecology of bat populations, although tolerance to anthropogenic activities has been described in some bat species (Jung & Threlfall, 2018; Russo & Ancillotto, 2015). Altogether, both historical and more contemporary factors can have significant implications for the long-term survival and conservation of island endemic bats.

Genetic analyses have become an essential tool for studying the ecology and evolution of island endemic bats. However, because of complex histories including allopatric divergence, colonization, and hybridization on islands, studies have highlighted the need for the rigorous use of both mitochondrial and nuclear microsatellite markers (Kuo et al., 2015). Indeed, these markers have different evolutionary timescales, permitting to assess historical and contemporary population structure, as well as different inheriting modes, widely used to assess sex-specific life-history traits (Pinzari et al., 2023; Taki et al., 2021). Maternally-inherited mitochondrial DNA (mtDNA) can trace colonization histories and past divergence, and provide estimates of female site fidelity and dispersal, while polymorphic nuclear microsatellite DNA are good candidates to infer recent gene flow and can provide information for both sexes.

By examining the genetic diversity and structure of island endemic bat populations, we can infer drivers of gene flow, estimate population sizes, and understand demographic history. For example, genetic analysis of the mastiff bat (*Molossus milleri*, family Molossidae) occurring on Jamaica, Cuba, and the Cayman Islands suggests that populations underwent bottlenecks, likely due to climate change in the early Pleistocene (Loureiro et al., 2020). Moreover, several studies have shown stronger genetic structure in the philopatric sex (mostly female), resulting from sex-biased dispersal behaviours in bat populations (Halczok et al., 2018; Jang et al., 2021; Moussy et al., 2013; Naidoo et al., 2016). In addition, bats often exhibit seasonal behaviours, in relation to change in food availability, habitat use, or reproductive cycle, and these factors may play critical roles in shaping genetic diversity patterns (Moussy et al., 2013). For example, in the little brown bat (*Myotis lucifugus*, family Vespertilionidae) and the northern long-eared bat (*M. septentrionalis*), individuals at swarming sites in autumn displayed a greater mtDNA genetic diversity than those at summering sites suggesting that swarming sites gather individuals from several summering sites (Johnson et al., 2015). Studies of genetic structure of island endemic bat species has mainly been carried out at the scale of multiple neighbouring islands (archipelago), but studies at a local scale, investigating sex or season variations, are still limited (Ratrimomanarivo et al., 2009). Obtaining a comprehensive picture of the local genetic structure of island endemic bats requires a fine-scale sampling scheme, including material from multiple sites and from different seasonal periods, thus allowing the detection of subtle diversity patterns, especially on islands of reduced size.

The Reunion free-tailed bat (*Mormopterus francoismoutoui*, family Molossidae) is a tropical insectivorous bat endemic to Reunion Island. This volcanic *in-situ* formed island is located in the southwestern Indian Ocean (Mascarene Archipelago) and emerged from the sea about 3 million years ago (Cadet, 1980). Reunion is located 950 km east of Madagascar, which is home to the Peter’s wrinkle lipped bat (*M. jugularis*), and only 175 km southwest of Mauritius Island, which is home to the Natal free-tailed bat (*M. acetabulosus*). Although small in size (2,512 km²), Reunion Island is shaped by a mountainous landscape, which could represent a barrier to bat dispersal, with the highest point at 3,070 m (Piton des Neiges) and a still active volcano (Piton de La Fournaise). *Mormopterus francoismoutoui* is broadly distributed on the island and roosts in different natural settings, such as caves and cliffs. This bat species had adapted to anthropogenic settings and thrives in the lowland urbanized areas where numerous roost sites occur in buildings and under bridges (Augros et al., 2015; Goodman et al., 2008). However, little is known in this species on how urbanization might modify life-history trait, population size and genetic structure. A recent longitudinal monitoring of several roosts revealed highly dynamic roosting behaviours (Aguillon et al., 2023). Specifically, large female aggregations (up to 50,000 pregnant individuals) within a limited number of maternity roosts are observed synchronously during austral summer, which coincides with a female-biased sex-ratio at the roost (Aguillon et al., 2023; Dietrich et al., 2015), and suggest female philopatry in this species. Moreover, towards the end of the austral summer, there is a decrease in roost size and a shift in sex-ratio (from female to male-biased), suggesting important seasonal sex-specific movements on the island. These details support the results of the first genetic study of this species, based on a limited number of samples (*n* = 31), that suggested little genetic structure and no isolation by distance within the island (Goodman et al., 2008).

In order to investigate the evolutionary history and genetic structure of *M. francoismoutoui*, we used an extensive spatio-temporal field sampling and the complementary information of microsatellite and D-loop mtDNA markers. We first analysed the evolutionary and demographic history of this species, by examining its relationship to other regional *Mormopterus* bats on neighbouring islands (Madagascar and Mauritius) and by testing the hypothesis of a recent population expansion linked to urbanization. We then analysed spatio-temporal patterns of genetic diversity and population structure across roosts all over Reunion Island and during different seasons. We specifically tested female philopatry and seasonal changes in the genetic structure linked to the dynamic roosting behaviour of this species. We expected that genetic structure to be more prominent in females during summer, and lower during the mating season because of mixing of individuals within roosts.

## Material and Methods

### Field sampling

Samples were collected at 18 roosts (coded with a 3-letter code) across Reunion Island (Fig. 1) and during different seasons. Among these roosts, six (AOM, CIT, PBV, RAC, STJ, and TM5) were only sampled once throughout the study, because of opportunistic sampling (Table S1). The remaining roosts were sampled multiple times from October 2018 to March 2020. Specifically, we collected samples from eight roosts (ESA, MON, PSR, RBL, RPQ, STM, TGI, and VSP) during three biological seasons: (i) the pregnancy period (austral summer) from late October to early December 2018), (ii) the non-reproductive period (austral winter) in June and July 2019, and (iii) the putative mating period in March 2019 and 2020 (Aguillon et al., 2023, Table S1). Three roosts (EGI, RES, and TBA) were quasi-empty during the non-reproductive/winter period, explaining the lack of data during this season. For one roost (TRI), no samples were collected in March because of logistic constraints.

**Figure 1.**
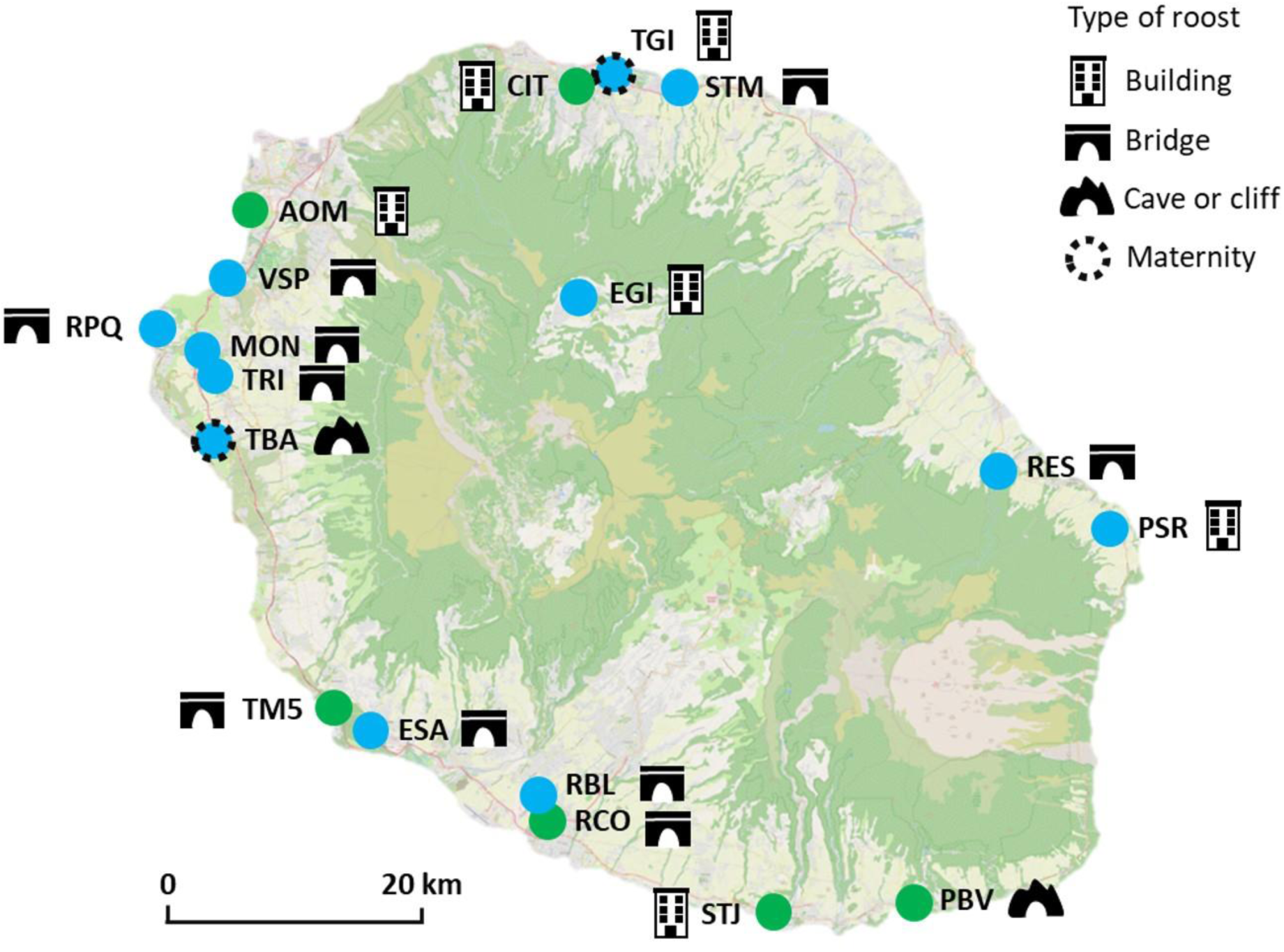
Sampling sites of *Mormopterus francoismoutoui* on Reunion Island. Details on roost sites are presented in Table S1. The twelve roosts in blue were monitored regularly over two years while the six in green were sampled only once. The green colour indicating forested areas and the pale blue indicating urbanized areas. Modified from Aguillon et al. (2023).

Bat captures took place during the dusk emergence, reaching a maximum of 60 individuals per night as described in Aguillon et al., (2023). We mainly used harp traps (Faunatech Ausbat) and Japanese monofilament mist nets (Ecotone) set close to the roost exit. Because of difficulties in installing harp traps or mist nets at the exit of some roosts, we sometimes employed a butterfly net on an elongated pole to catch bats, by carefully approaching resting individuals during the day. After capture, bats were immediately hydrated with water using a sterile syringe and placed in a clean individual bag close to a warm source (hot water bottle), and processed at the capture site. We visually ascertained the sex and age of each individual. Age was determined by examining the epiphysis fusion in finger articulations that are not ossified for juveniles. Wing punch samples (∼ 2 mm) were taken on each wing, stored in a cool box in the field before being transferred at -80°C at the laboratory. Finally, each bat was tattooed on the right propatagium with an individual alphanumeric code and released at the capture site.

Handling of bats was performed using personal protective equipment and gloves were disinfected between each individual bat and changed regularly, and all the equipment was disinfected between sites as well (see protocol in Aguillon et al., 2023 for more details). Bat capture and manipulation techniques were evaluated by the ethic committee of Reunion Island, approved by the Ministère de l’Enseignement Supérieur, de la Recherche et de l’Innovation (APAFIS#10140-2017030119531267), and conducted under a permit (DEAL/SEB/UBIO/2018-09) delivered by the Direction de l’Environnement, de l’Aménagement et du Logement (DEAL) of Reunion Island.

Samples (organ pool: spleen, lung, kidney) of *M. acetabulosus* from Mauritius in 2012 and *M. jugularis* from Madagascar (2012-2013) were obtained from vouchered individual batscollected during zoonotic disease studies (Gomard et al., 2016; Joffrin et al., 2020; Mélade et al., 2016), and from three different roosts on each island. Mauritius samples were collected under a memorandum of agreement for the supply of biological material by Government of Mauritius (delivered by the National Park and Conservation Service for authorization of Mauritius), signed on 17 December, 2010. Madagascar samples were collected under the permits delivered by the Direction du Système des Aires Protégées and Direction Générale de l’Environnement et des Forêts: no. 350/10/MEF/SG/DGF/DCB.SAP/SCB, no. 032/12/MEF/SG/DGF/DCB.SAP/SCBSE, no. 067/12/MEF/SG/DGF/DCB.SAP/SCBSE, no. 194/12/MEF/SG/DGF/DCB.SAP/SCB, no. 283/11/MEF/SG/DGF/DCB.SAP/SCB, no. 077/12/MEF/SG/DGF/DCB.SAP/SCBSE, no.238/14/MEEF/SG/ DGF/DCB.SAP/SCB, and no. 268/14/MEEF/SG/DGF/DCB.SAP/SCB.

### DNA extraction, PCR, sequencing, and genotyping

Wing punch samples of Reunion free-tailed bats were processed with the Cador Pathogen 96 Qiacube HT kit (Qiagen, Hilden, Germany). Samples were lysed before DNA extraction, in 180 μL of ATL buffer and 20 μL of Proteinase K at 56°C during 1h30. Then, the buffer VXL mixture was prepared replacing Proteinase K by sterile water. Total nucleic acids were extracted in an automated extractor Qiacube with slight modifications of the Q Protocol, including 350 μL of ACB, 100 μL of AVE, and 30 μL of TopElute. Nucleid acids from Mauritius and Madagascar samples were already available (protocols of extraction described in Gomard et al., 2016; Joffrin et al., 2020; Mélade et al., 2016).

Subsequently, a fragment of the D-loop region was amplified by PCR (expected: 896 pb) in a 20 μL reaction mixture containing 2 μL of DNA, 10 μL of GoTaq® Green Master Mix 2X (Promega, Madison, Wisconsin, United States), 1 μL of each primer at 10 μM D-loop-F (5’-CAAGACTTCAGGAAGAAGCTAACA-3’) and D-loop-R-Lg (5’-TATTCGTATGTATGTCCTGTAACCA-3’). PCR program included an initial denaturation step (95°C for 2 min), followed by 35 cycles of denaturation (95°C for 30 sec), annealing (50°C for 30 sec), elongation (72°C for 1 min 30 sec), and a final elongation step (72°C for 7 min). PCR products were Sanger-sequenced by the GENOSCREEN platform (Lille, France). The D-loop chromatograms were visually checked using Geneious 9.1.8 (Biomatters Ltd, Auckland, New Zealand) and sequences were aligned using CLC Sequence Viewer 7.6.1 (Qiagen Aarhus A/S, Aarhus, Denmark). DNA extracted were genotyped by GENOSCREEN, using a panel of 12 previously described microsatellite markers, according to the protocol and primers of Dietrich et al. (2019).

### *Analyses of* M. francoismoutoui

Using mitochondrial D-loop data, genetic diversity indices, including haplotype number, haplotype diversity (Hd), and nucleotide diversity (π), were measured at the roost-level using DnaSP v6.12.03 (Rozas et al., 2017), for the entire Reunion Island dataset and then separately for each sex and season. Differences among roosts, sexes, and seasons were tested using analyses of variance (ANOVA) in RStudio 1.4.1106 (RStudioTeam, 2021). Spatial structure was assessed by calculating genetic distances (Φ^st^) among roosts for the entire Reunion Island dataset and then separately for each sex and season using Arlequin 3.5.2.2 (Excoffier & Lischer, 2015). The significance of multiple tests was corrected with the Holm method using RStudio. To test for temporal differences in the spatial structure, Φ_st_ values were compared among seasons using ANOVAs and Tukey’s post-hoc tests. Subsequently, to test for the presence of isolation by distance (IBD), we performed a Mantel test with 1,000 permutations using Arlequin and calculated the correlation between genetic and geographic distances. IBD was first tested for the whole Reunion Island dataset, and then separately for each sex and season. Finally, we also used an AMOVA test (analysis of molecular variance) in Arlequin and defined “population” as individuals from a single roost to test for a roost-associated genetic structure in the Reunion population. The significance of this test was assessed by 1,000 permutations of individuals among roosts.

We determined the most appropriate nucleotide substitution model of Reunion Island D-loop sequences based on AIC criterion (Akaike, 1974) using JModelTest v2.1.10 (Darriba & Posada, 2016). We constructed a Bayesian tree using BEAST v.2.6.4 (Bouckaert et al., 2019) with TN93 site model with invariant and gamma distribution (Tamura & Nei, 1993) including roost location as a trait. We used an uncorrelated lognormal relaxed molecular clock (Drummond et al., 2006) of 0.2 substitutions/site/million years (Petit et al., 1999), with a 100 million chain length and sampling every 10^4^ steps, and a burning of 10%. We ran three analyses and combined log outputs (removing 10% of burning for each output) using LogCombiner v2.6.4 (Rambaut & Drummond, 2015). Traces of Markov Chain Monte Carlo (MCMC) were checked for convergence of the posterior estimates of the effective sample size (ESS) to the likelihood using Tracer v1.7.1 (Rambaut et al., 2018). We combined tree outputs (removing 10% of burning for each output) to obtain a consensus tree using LogCombiner v2.6.4 (Rambaut and Drummond, 2015) and then TreeAnnotator v2.6.4 (Rambaut & Drummond, 2019).

To investigate the demographic history of *M. francoismoutoui*, we calculated the expected frequency distributions of pairwise differences between D-loop sequences (mismatch distribution) in DnaSP v6.12.03 (Rozas et al., 2017). We also used a neutrality test with 1,000 simulated samples using Arlequin v.3.5.2.2 (Excoffier & Lischer, 2015) based on Harpending’s raggedness index (r, Harpending, 1994), Fu’s Fs (Fu, 1997), and the sum of squared deviations (SSD) between observed and expected mismatch indices. We then calculated the expansion time using the formula 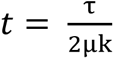 from Rogers & Harpending (1992) where τ is the expansion date calculated with the mismatch distribution, μ is the mutation rate, and k is the average number of nucleotide sites per haplotype. Global population size change through time was reconstructed using a coalescent Bayesian skyline model (CBS, Drummond et al., 2005) with the same parameters as described above in BEAST v.2.6.4 (Bouckaert et al., 2019). We performed BEAST analyses changing the dimension group parameter from 3 to 10 groups, according to the results of the Yule Bayesian tree. We ran three analyses for each group and followed the same method described for Yule Bayesian tree and choose the best k according to the higher ESS.

For the microsatellite data, genotype determination was performed using GeneMapper 6 (ThermoFisher). We tested the dataset for scoring errors, out of range allele, and null alleles for each roost using Microchecker v.2.2.3 (Van Oosterhout et al., 2004). Using GenAlex 6.503 (Smouse & Peakall, 2012), the global number of genotypes was calculated, and for each roost, deviations from Hardy-Weinberg equilibrium were tested for each locus. A permutation test with 1,000 permutations was used to perform linkage disequilibrium analysis between each pair of loci using Genetix v.4.05.2 (Belkhir et al., 2004). We used Fstat v.2.9.4 (Goudet, 2003) to calculate the inbreeding coefficient (F_IS_) within each roost (Weir & Cockerham, 1984) and tested the significance by randomizing alleles among individuals within roosts (5,000 permutations). We estimated the observed (Ho) and expected (He) heterozygosity in each roost, first using the whole dataset, subsequently estimated these two parameters for each sex and season, and tested for sex and temporal differences using an ANOVA in RStudio. Effective population size (N_e_) was estimated for the global population because of a lack of genetic structure (see results) with the linkage-disequilibrium model and assuming random mating using NeEstimator v2.1 (Do et al., 2013). For this, we used a minimum allele frequency of 0.05 and 0.02 to calculate upper and lower limits of N_e_.

Genetic distances (F_st_) between roosts were calculated globally, and then separately for each sex and season, using Arlequin 3.5.2.2 (Excoffier & Lischer, 2015). The significance of multiple tests was corrected with Holm method using RStudio. To test for temporal differences in the spatial structure, F_st_ values were compared among seasons using ANOVAs and Tukey’s post-hoc tests. To check for IBD, we performed Mantel tests (1,000 permutations) using Arlequin 3.5.2.2 (Excoffier & Lischer, 2015), as for mitochondrial data.

To test for a roost-associated genetic structure in the population, we used an AMOVA in Arlequin, as described for mtDNA. Then, we employed STRUCTURE v2.3 (Pritchard et al., 2010), and performed two analyses with and without the LocPrior model, which uses location to test for a weak signal of population structure (Hubisz et al., 2009). We used the admixture model with correlated allele frequencies among groups, and 10 replicate runs were performed with a burn-in of 10^6^ steps and 10^6^ recorded steps for the Monte Carlo Markov Chain (MCMC). We ran K from 1 to 19 groups (corresponding to the number of studied roosts, plus one). We applied the Evanno method (Evanno et al., 2005) to estimate the best K, but results were inconclusive (see results). To determine the optimal number of genetic clusters, we performed a k-means clustering analysis, tested K from 2 to 19 over 26 indices according to the “majority rule”, using the *NbClust* package in Rstudio. We also performed a principal coordinate analysis (PCoA) using GenAlex 6.503 to visualize possible genetic clusters.

Finally, to compare mitochondrial and nuclear results, we overlaid the genetic clusters identified using microsatellite markers and clades depicted by the BEAST phylogeny obtained with the D-loop sequences. Based on the dimension group parameter estimated by the best skyline converging model in BEAST, we defined five mtDNA clusters according to posterior probabilities > 0.99 (see results). We used a generalized linear model (GLM) in Rstudio, including the nuclear clusters as the numeric response variable (value of PC1 from PCoA) and the mtDNA genetic clusters as the explanatory response.

### *Genetic relationships among regional* Mormopterus

In order, to investigate genetic relationships among the three regional *Mormopterus* species, we reconstructed a Bayesian tree based on Yule model (Yule, 1925), using the island as a trait to resolve the spatial origin of the nodes using BEAST v.2.6.4 (Bouckaert et al., 2019). HKY model (Hasegawa et al., 1985) with invariant and gamma distribution was used according to the best substitution model on AIC criterion using JModelTest (Darriba & Posada, 2016). We used the same parameters in BEAST as described above for the Reunion Island dataset.

Microsatellite analysis of regional samples was performed using STRUCTURE v2.3 (Pritchard et al., 2010), with LocPrior (Hubisz et al., 2009) and no admixture model, using uncorrelated allele frequencies among group. We ran 10 replicates with a burn-in of 10^6^ steps, 10^6^ recorded steps for the MCMC, and K from 1 to 5 groups (corresponding to the number of species, plus two). We apply the Evanno method (Evanno et al., 2005) to estimate the best K.

## Results

### Data quality

Altogether, we obtained good quality D-loop sequences for 603 *M*. *francoismoutoui* (alignment of 985 pb) and 30 sequences for each *Mormopterus* species (Mauritius and Madagascar, 811 pb). We genotyped 1,136 individuals of *M*. *francoismoutoui* using the 12 microsatellite loci, and 30 individuals from Mauritius (*M*. *acetabulosus*) and Madagascar (*M*. *jugularis*). One locus (MF_loc11) was removed because of a high percentage of uninterpretable weak signals in the Reunion Island dataset (∼ 41% of individuals). Also, we removed individuals from Reunion for which at least six loci were not genotyped (*n* = 22), leading to a final microsatellite data set containing 3.7% of missing alleles. In the Reunion dataset, a majority of loci significantly deviated from Hardy-Weinberg equilibrium, especially MF_Loc03, MF_Loc04, MF_Loc05, and MF_Loc15. Null alleles were detected in several loci, especially in MF_Loc04, and MF_Loc015. Three loci were implicated in several linkage disequilibria: MF_Loc03, MF_Loc05, and MF_Loc28.

### *Genetic relationship among regional* Mormopterus

The time calibrated Bayesian phylogeny strongly supported three genetic clades corresponding to each bat species, thus confirming their monophyly (Fig. 2). Interestingly, STRUCTURE analyses identified two genetic clusters, separating the Malagasy species in one cluster, and the two species on Reunion and Mauritius in a second cluster (Fig. S1). Surprisingly, when K = 3, only 20% of the runs assigned each bat species to a different clusters. Indeed, most of the runs for K = 3 (80%) grouped Reunion and Mauritius bats in the same genetic cluster, while Malagasy bats were composed of two clusters. The BEAST analysis showed that the inferred TMRCAs for the three *Mormopterus* species was 374,800 years ago (HPD: 287,500 – 467,000) and the divergence of *M*. *francoismoutoui* on Reunion from *M*. *acetabulosus* in Mauritius was dated at 278,000 years ago (HPD: 209,100 – 355,100). The diversification of *M*. *jugularis* on Madagascar was dated at 255,200 years ago (HPD: 192,800 – 319,200) and occurred later for the species on Mauritius (181,600; HPD: 135,800 – 232,500) and Reunion (171,400; HPD: 129,600 – 218,000).

**Figure 2.**
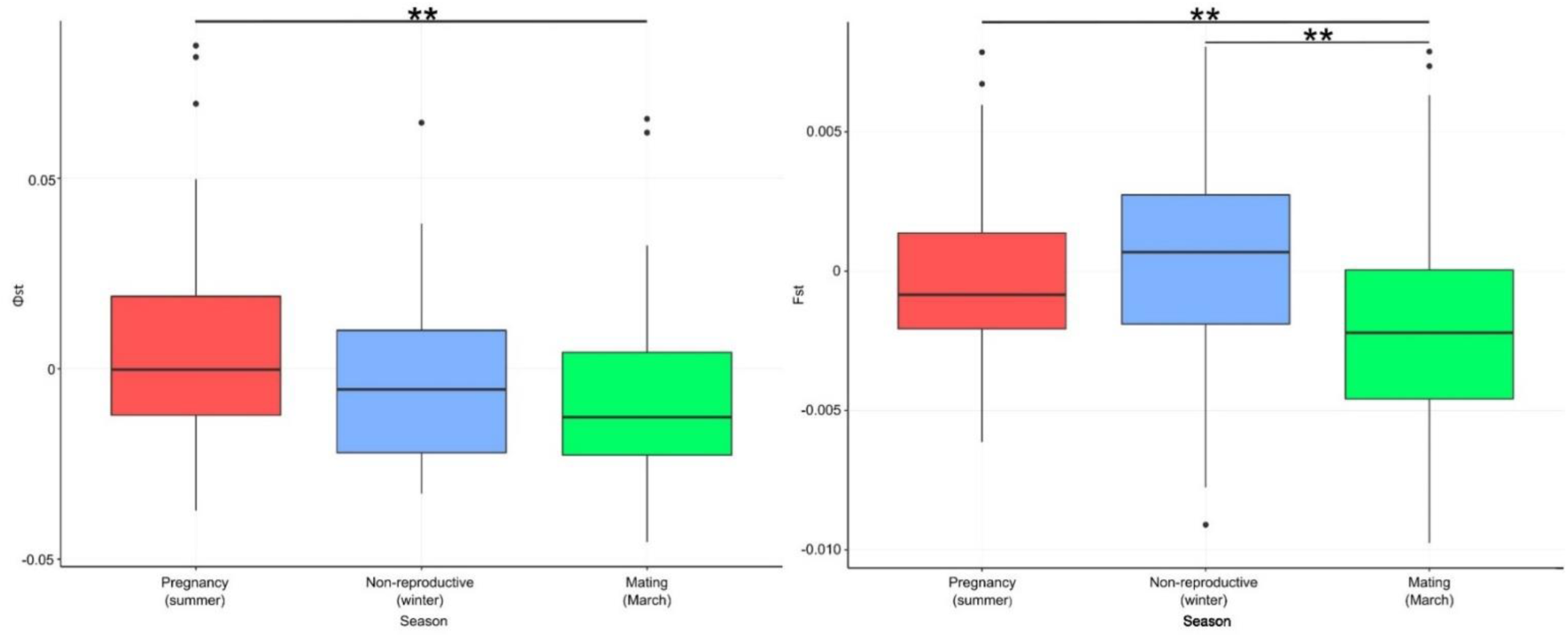
Temporal differences in Φ_st_ (mtDNA) and Fst (microsatellites) values between seasons in *Mormopterus francoismoutoui*. ** *p* < 0.01 (Tuckey’s post-hoc tests).

### *Genetic diversity of* M. francoismoutoui

Mitochondrial DNA and microsatellite markers revealed a high genetic diversity within the population of *M*. *francoismoutoui*. For mtDNA, 410 haplotypes (out of 603 sequences) were identified with an average haplotype diversity (Hd) of 0.998 (Table 1) and a global nucleotide diversity (π) of 0.0284. Using the microsatellite markers, we identified 1,135 genotypes (in 1,136 individuals), and only two individuals (both captured in the TGI roost) shared the same genotype. The average observed (Ho) and expected (He) heterozygosities were high (Ho = 0.778 ± 0.009, He = 0.797 ± 0.008) and there was no evidence of inbreeding between individuals occupying the same roosts, as none of the F_IS_ values were significantly different from zero (Table 1). For both the mtDNA (Hd and π) and nuclear (Ho and He) data, there was no significant differences in the level of genetic diversity between roosts, nor between sexes and seasons (ANOVA, all *p* > 0.05, Table S2 and S3 for D-loop, Table S4 and S5 for microsatellites).

**Table 1.** Global genetic diversity indices of *Mormopterus francoismoutoui*, calculated with mitochondrial DNA (D-loop) and 11 microsatellite markers. For roosts details see Figure 1 and Table S1. N: number of individuals, Hd: haplotype diversity, π: nucleotide diversity, Ho: observed heterozygosity (± standard error), He: expected heterozygosity (± standard error), and Fis: inbreeding coefficient (none are significantly different from zero).

| Roost | Mitochondrial data |  |  | Microsatellite data |  |  |  |
| --- | --- | --- | --- | --- | --- | --- | --- |
| | N | Hd | $\pi$ | N | Ho | He | Fis |
| AOM | 14 | 0.989 | 0.0264 | 30 | 0.762 ± 0.035 | 0.793 ± 0.037 | 0.058 |
| CIT | 15 | 1.000 | 0.0304 | 29 | 0.787 ± 0.035 | 0.791 ± 0.029 | 0.022 |
| EGI | 41 | 0.999 | 0.0287 | 64 | 0.766 ± 0.040 | 0.807 ± 0.030 | 0.060 |
| ESA | 46 | 0.998 | 0.0272 | 96 | 0.776 ± 0.033 | 0.806 ± 0.033 | 0.044 |
| MON | 48 | 1.000 | 0.0291 | 88 | 0.797 ± 0.031 | 0.800 ± 0.035 | 0.009 |
| PBV | 11 | 1.000 | 0.0342 | 11 | 0.759 ± 0.032 | 0.762 ± 0.030 | 0.053 |
| PSR | 47 | 0.997 | 0.0297 | 96 | 0.786 ± 0.037 | 0.797 ± 0.036 | 0.018 |
| RAC | 16 | 1.000 | 0.0309 | 14 | 0.836 ± 0.070 | 0.772 ± 0.036 | -0.045 |
| RBL | 46 | 0.999 | 0.0282 | 93 | 0.763 ± 0.028 | 0.802 ± 0.035 | 0.055 |
| RES | 30 | 1.000 | 0.0284 | 62 | 0.744 ± 0.040 | 0.802 ± 0.033 | 0.081 |
| RPQ | 45 | 0.997 | 0.0297 | 100 | 0.788 ± 0.031 | 0.805 ± 0.034 | 0.026 |
| STJ | 15 | 1.000 | 0.0278 | 30 | 0.782 ± 0.029 | 0.796 ± 0.034 | 0.036 |
| STM | 43 | 1.000 | 0.0289 | 70 | 0.797 ± 0.037 | 0.809 ± 0.032 | 0.022 |
| TBA | 46 | 0.998 | 0.0290 | 63 | 0.783 ± 0.040 | 0.801 ± 0.034 | 0.032 |
| TGI | 49 | 0.997 | 0.0308 | 103 | 0.766 ± 0.044 | 0.806 ± 0.035 | 0.054 |
| TM5 | 15 | 1.000 | 0.0305 | 28 | 0.752 ± 0.044 | 0.791 ± 0.035 | 0.070 |
| TRI | 30 | 1.000 | 0.0299 | 58 | 0.781 ± 0.038 | 0.801 ± 0.031 | 0.034 |
| VSP | 46 | 0.999 | 0.0290 | 101 | 0.779 ± 0.035 | 0.804 ± 0.034 | 0.036 |
| TOTAL | 603 | 0.998 | 0.0284 | 1136 | 0.778 ± 0.009 | 0.797 ± 0.008 | 0.034 |

### Genetic structure within the Reunion Island population

Globally, for both mitochondrial and nuclear markers, no isolation by distance (r_mtDNA_ = 0.006, *p* = 0.46; r_nuclear_ = 0.11, *p* = 0.14) was detected nor significant pairwise differentiation among roosts (Fst and Φst, Table S6). Results were unchanged when analyses were performed separately for each sex and season (Table S7 for IBD results). However, we identified differences in Φst and Fst values among seasons (ANOVA, *p* = 0.003 and *p* = 0.001 respectively). The mitochondrial marker showed higher Φst values in the pregnancy period compared to those of the mating period (*p* = 0.002), while for microsatellites, Fst values during both the pregnancy (*p* = 0.008) and the non-reproductive period (*p* = 0.003) were higher compared to those during the mating period (Fig. 3). AMOVA results with both markers showed that genetic variation (100.40% for mtDNA and 99.99% for microsatellites) was largely due to differences among individuals within roosts. Results from the STRUCTURE analyses revealed no genetic clustering (no conclusive Evanno result, Fig. S2), while the k-means clustering analysis evaluated the optimal number to three genetic clusters (Fig. S3). This result was corroborated by the PCoA but with only a small genetic variation explained by the two first axes (PC1: 4.73% and PC2: 3.31%, Fig. 4). Interestingly, these three clusters included bats from the different roosts.

**Figure 3.**
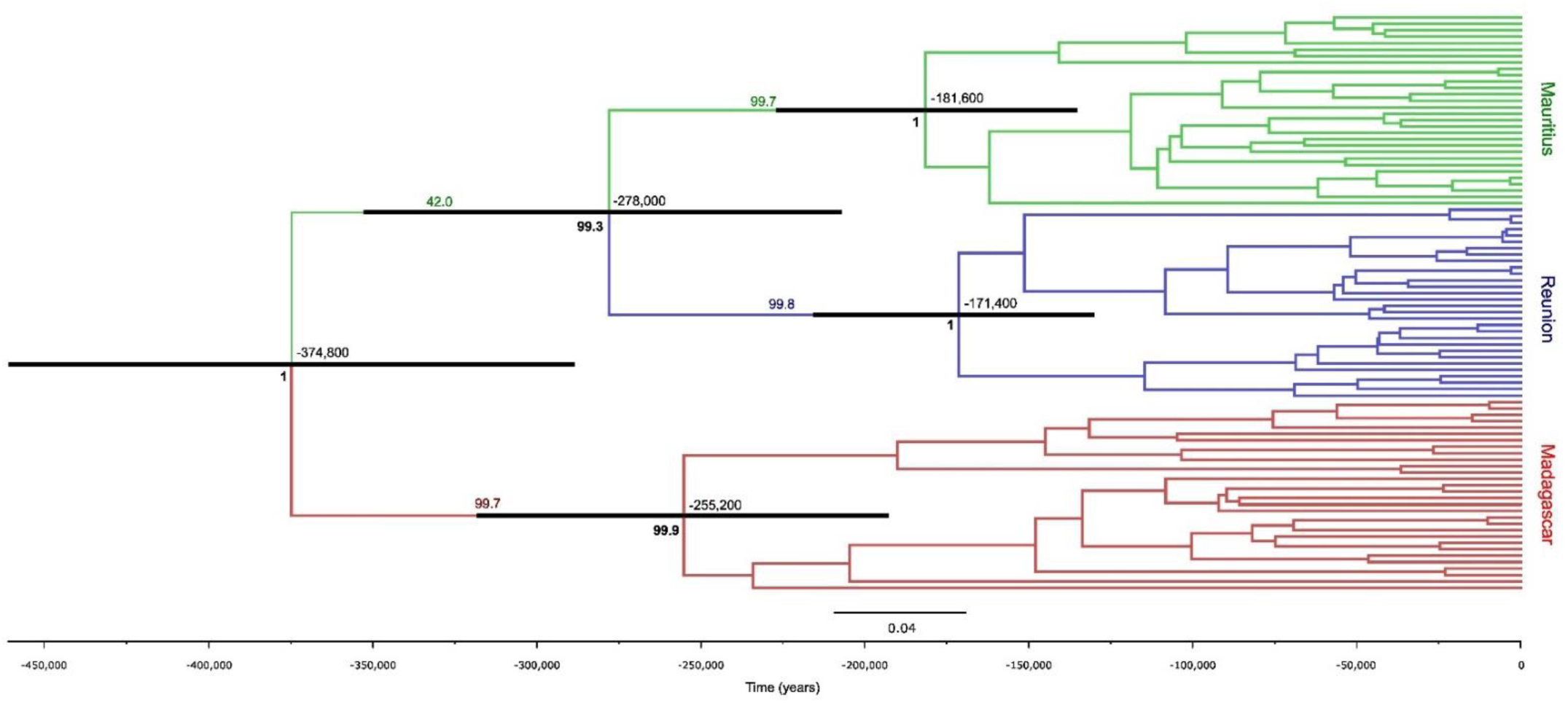
Bayesian tree topology inferred from mitochondrial data (D-loop) for three *Mormopterus* species occurring on southwestern Indian Ocean islands. The Yule model was used with HKY (I+G) substitution model and relaxed molecular clock of 0.2 substitution/site/million years. The time is indicated in the x-axis from past (left) to recent (right) time. Islands are colour coded: Mauritius (*M. acetabulosus*) in green, Reunion (*M. francoismoutoui*) in blue, and Madagascar (*M. jugularis*) in red. Branches are coloured according to the main location posterior probabilities. Posterior probabilities values are indicated in bold and the black horizontal bars represent the 95% HPD of node ages with main age values indicated above the node.

**Figure 4.**
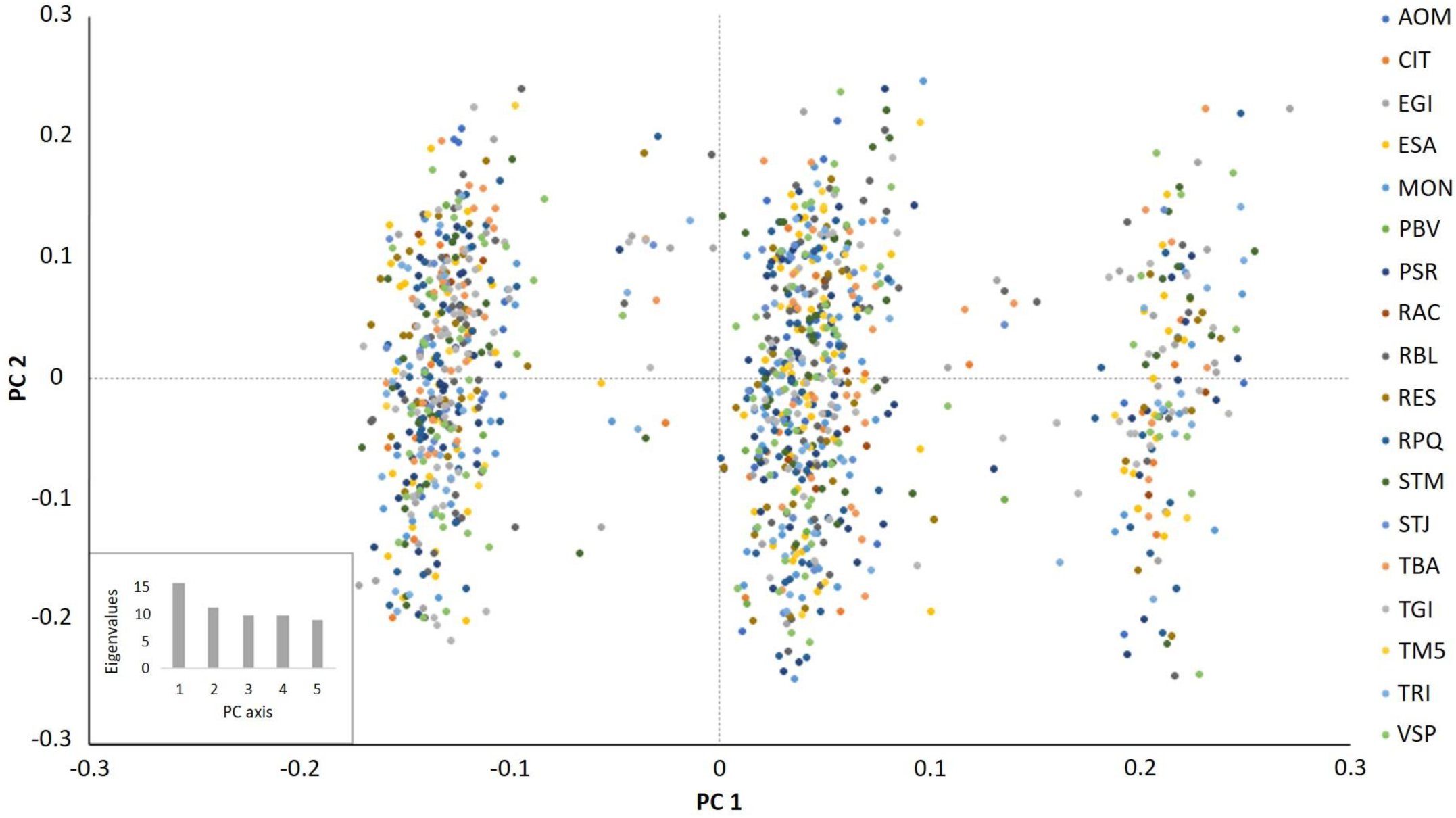
Principal Coordinates Analysis (PCoA) of microsatellite markers for *Mormopterus francoismoutoui*. Roosts are indicated by colours and the eigenvalues of the first five principal component (PC) are shown.

Based on a conservative posterior probability > 0.95, the Bayesian tree built with the Yule model of speciation showed at least three well-supported genetic clusters (Fig. 5). These clusters included bats from all roosts, but the best reconstruction of ancestral nodes failed to predict the roost origin of individuals. The best skyline converging model indicated the occurrence of five genetic groups (Fig. 5), followed by models with six and 10 groups with a close likelihood ESS (Table S8). These five clusters were separated by a maximum of 3.3% of divergence. When overlaying the genetic clusters identified with microsatellite markers on the BEAST phylogeny, no significant results were found revealing that genetic clusters from both markers are different (GLM, χ²_4_ = 0.03, *p* = 0.72).

**Figure 5.**
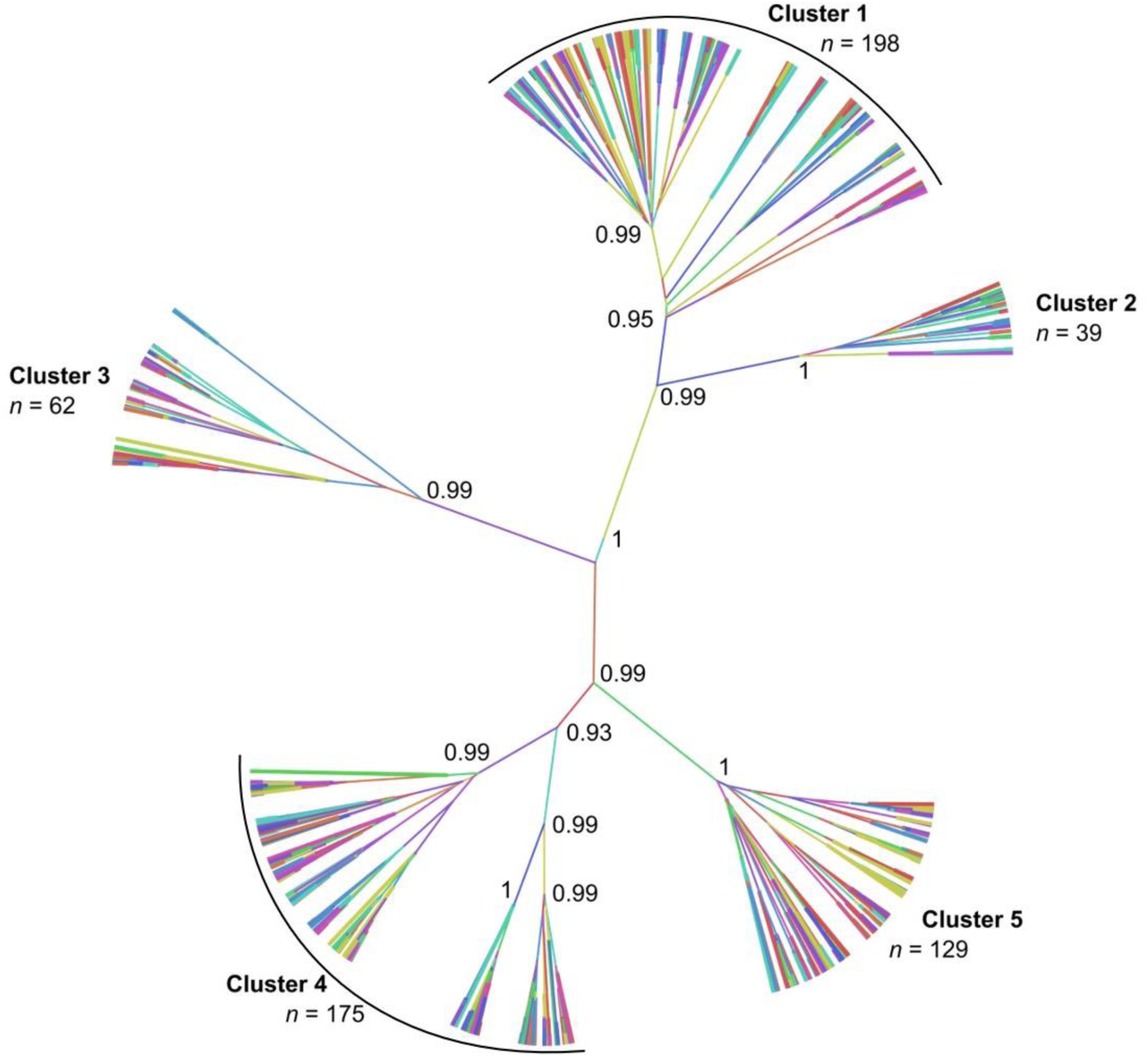
Bayesian tree topology inferred from mitochondrial data (D-loop) for the 18 roosts of *Mormopterus francoismoutoui*. The Yule speciation model was used with TN93 (I+G) substitution model and a relaxed molecular clock of 0.2 substitution/site/million years. Colours correspond to roost sites, and line weigh represents the probability of roost location. Posterior probability values higher than 0.95 are indicated next to the node of main genetic groups. Bold numbers correspond to genetic clusters based on the best skyline converging model and defined according to posterior probabilities > 0.99.

### *Population demographic history of* M. francoismoutoui

Estimations of effective genetic population size with the lowest allele frequency at 0.05 and 0.02 led to infinite estimate of Ne (95% CI_0.05_: 7783.5 - Infinite and 95% CI_0.02_: 22347.5 – Infinite). Moreover, the mismatch distribution under the expansion model showed a clear signal of demographic expansion with a multimodal distribution with three peaks (Fig. 6A). Raggedness index and Sum of Square Deviation had non-significant values under the model of demographic expansion (r = 0.0005, *p* = 1; SSD = 0.002, *p* = 0.71). The result of Fu’s Fs showed significant negative values indicating an excess of rare haplotypes compared to expected values under neutral model (Fs = -23.33, *p* = 0.04). All these results suggested an ancient demographic expansion in the population. Based on information calculated from the mismatch distribution test, we determined the expansion time 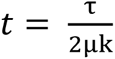 with τ = 29.156, μ = 0.2 substitutions/site/million years (Petit et al., 1999) and k = 811 pb. We found an expansion time *t* = 89,876 years. This was coherent with the Bayesian skyline plot showing a stable population size starting from 175,000 years and up to 90,000 years. Then, a slight increase in population size began and was followed by a drastic expansion around 55,000 years, lasting about 10,000 years. In the last 45,000 years, the population size still increased but at a slower rate. The end of the curve suggested a recent stabilization or a decrease in population size occurring about 500 years ago (Fig. 6B).

**Figure 6.**
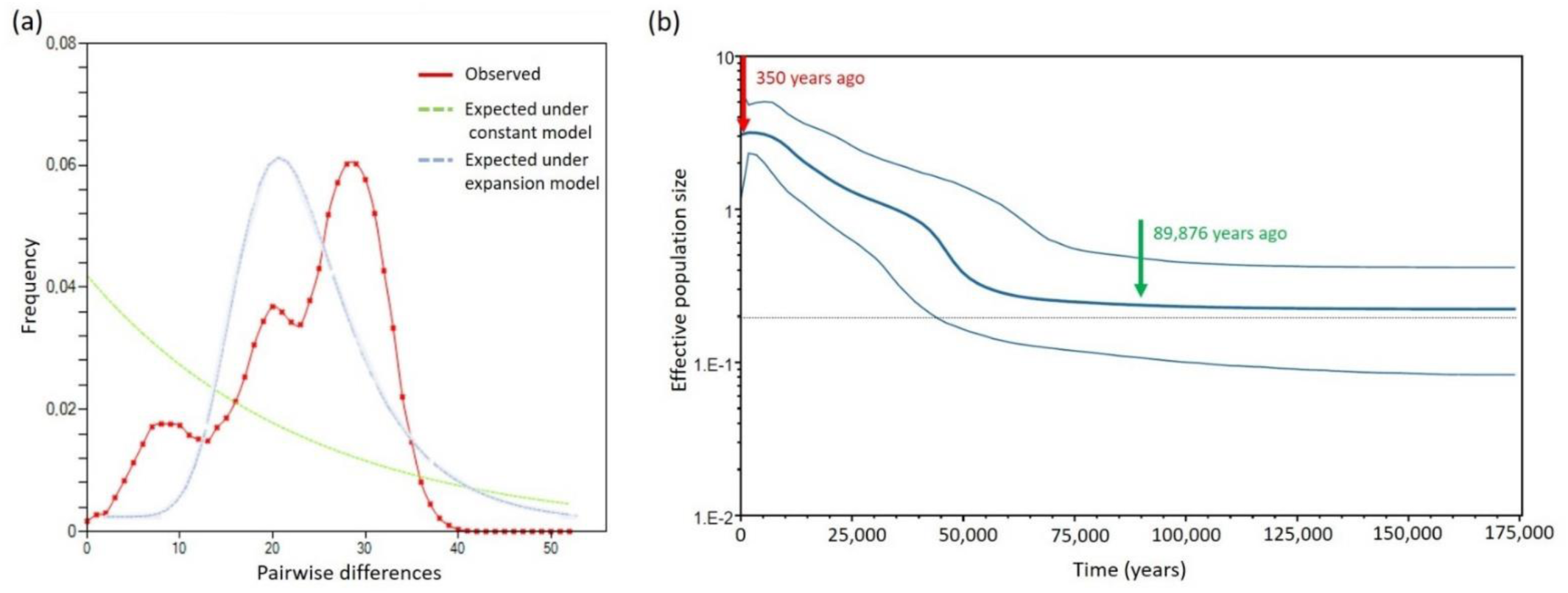
(a). Frequency of pairwise differences distribution (mismatch distribution) for *Mormopterus francoismoutoui*. The frequency observed is represented by the red dotted line, the expected frequency under the hypothesis of population constant model is indicated by the green line, and population expansion model by the blue line. **(b). Coalescent Bayesian skyline plot inferred from mitochondrial data (D-loop) showing estimated demographic history of *Mormopterus francoismoutoui* (with dimension group parameter = 5)**. Time is indicated in the x-axis from recent (left) to past (right) and the estimate effective population size (Ne) is represented in the y-axis. The central blue line is the median surrounded by the upper and lower estimates of 95% credibility interval. The Reunion free-tailed bat estimated time of expansion is indicated with the green arrow around 90,000 years before present and the red arrow represent the colonization of the island by humans since 350 years.

## Discussion

Despite living on the small oceanic island of Reunion (2,512 km²), our study revealed an extreme high genetic diversity in *Mormopterus francoismoutoui*, with 68% of unique D-loop haplotypes and 99.9% of unique microsatellite genotypes. Only one microsatellite genotype was found to be shared by two individuals, which were both captured in the same roost (TGI), suggesting a kinship link between them. Our results support those of Goodman et al. (2008) and are similar to previous studies reported in much bigger islands, such as in *M*. *jugularis* on Madagascar (587,041 km², Ratrimomanarivo et al., 2009) and *Myotis punicus* on the Mediterranean islands of Corsica (8,722 km²) and Sardinia (24,090 km², Biollaz et al., 2010). High levels of genetic diversity in island endemic bats can be explained by large population size (Frankham, 1996), which is supported by our results providing infinite large effective population size estimates for *Mormopterus francoismoutoui*. Such results are coherent with the fact that Molossidae bats form large and dense colonies (e.g. *Tadarida brasiliensis* of the family Molossidae, McCracken & Wilkinson, 2000) and corroborate our field observations of numerous roosts across Reunion Island, with an estimated current population size probably far over 120,000 individuals (Aguillon et al., 2023).

Our Bayesian phylogenetic analysis indicates that *M. francoismoutoui* form a distinct monophyletic lineage that diverged about 278,000 years ago from *M*. *acetabulosus*. The monophyly of the Reunion species supports the hypothesis of a single colonization event by overwater dispersal, although the geographic origin of its ancestor could not be determined in our study. A previous taxonomic work on *Mormopterus* bats from the Mascarene Islands showed morphological similarities with *M*. *norfolkensis* from Australia, while *M*. *jugularis* from Madagascar was reported closer to *M*. *doriae* of Sumatra (Goodman et al., 2008; Peterson, 1985). The different patterns of grouping among southwestern Indian Ocean islands for members of the genus obtained with STRUCTURE (Fig. S1) may indeed suggest different geographic origins for Mascarene and Madagascar *Mormopterus*, respectively. Although detailed genetic studies are available for certain regions of distribution of taxa currently placed in the genus, such as here for the southwestern Indian Ocean islands and Reardon et al. (2014) for the Australian species, no broad analysis is available across the broad geographic range of members of the genus. This will be needed to better define the origin of the southwestern Indian Ocean island species.

Once established on Reunion Island, the population of *M*. *francoismoutoui* remained stable and started increasing slowly 90,000 years ago, with a remarkable expansion around 55,000 years ago (Upper Pleistocene). This timing coincides with the estimated period when both volcanos (Piton des Neiges and Piton de la Fournaise) were active: from 65,000 years to 20,000 years (Nehlig & Marie, 2005). Such volcanic activity could have created new suitable habitats (like cliff crevices and caves), and thus enhanced range expansion of this species. Indeed, it has been previously suggested that bat species may benefit from volcano activity like the New Zealand short tailed bat (*Mystacina tuberculate*, family Mystacinidae) that experienced a range expansion possibly following rapid reforestation after a volcanic eruption (Lloyd, 2003).

Reunion Island was colonized by humans 350 years ago, with a drastic human population expansion during the 20^th^ century (Sandron, 2007). Contrary to our predictions that recent urbanization of Reunion Island, specifically construction of permanent structures (i. e. buildings, bridges) might have enhanced bat population size, our results suggest that the population expansion of *M*. *francoismoutoui* stopped (or slowed down) about 500 years ago. This result could be due to model error and based on a single locus (Ho & Shapiro, 2011). However, such a pattern of human intervention and increased population size has already been described associated with 17^th^ century deforestation for several Amazonian bats of the family Phyllostomidae (Silva et al., 2020). After human colonization of Reunion Island, the ecosystems were profoundly and rapidly modified (Lagabrielle et al., 2009). *Mormopterus francoismoutoui*, which is not a forest-dwelling species, now use for roosting sites some of the few remaining relatively large caves, as well as day-roosting sites in urban areas. Given the odour of the bats occupying roost sites in human constructions, numerous urban roosts are discouraged and a by-product of this might be higher rates of bat mortality (Augros et al., 2015). Moreover, extensive landscape modifications and human activities may have changed behaviour and physiology of this species, and may negatively affect the fitness of individuals and population size, as has been shown for other bat taxa (Russo & Ancillotto, 2015). The evolution of *M*. *francoismoutoui* population size over recent decades has not been assessed. The large and dense populations at roost sites of this species make the direct counts of individual bats difficult, occupancy modelling based on acoustic survey data, together with data from recaptured bats and mark-recapture models, should provide an alternative method to precisely assess trends in population size in relation to human activities (Oyler-McCance et al., 2017; Rivers et al., 2006; Rodhouse et al., 2019).

Our results suggest that the large current population of *M*. *francoismoutoui* experiences important levels of gene flow, as no significant genetic differentiation among roosts was found, as well as, low levels of inbreeding and no isolation by distance across sampled populations. Further, the high genetic diversity linked to the large population size may counter-balance a weak signal of spatial genetic structure (Gauffre et al., 2008). Interestingly, we found stronger Φst values during the summer, which might be associated with some degree of female philopatry during the pregnancy and parturition periods and supported with the massive aggregations of pregnant females observed during this period, specifically at the TBA roost (Table S1, Aguillon et al., 2023; Dietrich et al., 2015). Subsequently, in March, Φst values were the lowest, suggesting that bats dispersed and mixed within roosts coherent with the mating period (Moussy et al., 2013). Interestingly, the Fst values calculated with the microsatellites were still high during the non-reproductive winter months. This discordance between markers may be the consequence of behavioural aspects of males, which are probably more sedentary during non-reproductive period, and fits with observations that bats, particularly adult females, leave the studied roost sites during winter and disperse to unknown wintering sites (Aguillon et al., 2023). Our results thus suggest high levels of dispersal in this species across Reunion Island, and its capacity to disperse over the island’s mountainous landscape. However, it is important to note a similar lack of genetic structure in *M*. *jugularis* on Madagascar and not related to sex classes (Ratrimomanarivo et al., 2009). These results likely indicate a common evolutionary trend among *Mormopterus* species on southwestern Indian Ocean Islands, and more broadly in Molossidae species, such as the Mexican free-tailed bat *Tadarida brasiliensis* (Glass, 1982; McCracken et al., 2008) capable of long-distance migration and high altitude displacements. Interestingly, roosts occupied by *M*. *francoismoutoui* are often located in urban areas which could have facilitated dispersal by increasing connectivity between populations as molossid bats in general seem less affected by human-induced land changes (Richardson et al., 2021; Russo & Ancillotto, 2015).

Despite the absence of spatial genetic structure, our phylogenetic analyses showed that *M*. *francoismoutoui* has diversified into at least five deeply divergent mtDNA lineages, that are found in sympatry at the level of roost sites, including maternity roosts. Sympatric mtDNA lineages are not commonly described in animals, and especially in bats (Andriollo et al., 2015; Sun et al., 2016) and their origin and maintenance are often difficult to resolve (Hogner et al., 2012; Makhov et al., 2021; Webb et al., 2011). This may be explained by stochastic lineage sorting processes that occur in panmictic populations with large effective population size (Hogner et al., 2012; Webb et al., 2011). Also, we cannot exclude the possibility of female philopatry that would increase population structure in the mtDNA marker (Moussy et al., 2013), and this remained the same even when genetic analyses were performed for each sex. The divergence of mtDNA lineages may also reflect long periods of geographical isolation after the colonization of Reunion Island by the ancestral population. Our Bayesian phylogeny dated the start of the *in-situ* diversification back to 175,000 years ago, and the sharp increase in population size (about 55,000 years ago) coincides with the apparition of multiples lineages within the population (Fig. 2). Interestingly, the mtDNA genetic structure was not observed in the microsatellite analyses. Although increased allelic homoplasy at microsatellite loci may mask genetic differentiation over long periods of time in species with large populations (Estoup et al., 2002), our results may also suggest that the divergent mtDNA lineages are not reproductively isolated and that recent admixture of ancient lineages might have contributed to the high nuclear polymorphism detected (Andriollo et al., 2015; Sun et al., 2016). Indeed, such recent gene flow would erase genetic signatures at microsatellite loci more rapidly than mtDNA loci, explaining the absence of a strong signal of nuclear structure.

However, in the case of recent gene flow in *M*. *francoismoutoui*, our nuclear data suggest that it does not occur randomly within the population, as the clustering analysis and PCoA on microsatellite markers revealed the presence of three distinct clusters (also in sympatry within roosts). More interestingly, these nuclear clusters did not overlay mtDNA clusters and were not detected with the STRUCTURE analysis. This discordance between both markers can be explained by different evolutionary time processes and inheritance (Harrison, 1989; Toews & Brelsford, 2012). Such opposite patterns between markers have previously been described in different bat species (Laine et al., 2023; Naidoo et al., 2016; Sun et al., 2016) and highlight the need to use several markers for reconstructing complex evolutionary histories (Kuo et al., 2015). The microsatellite structure may correspond to a few isolated mating roosts on the island or could reflect putative adaptations like morphological or acoustic differences, as described for the big-eared horseshoe bat (*Rhinolophus macrotis*, family Rhinolophidae, Sun et al., 2016). Further bat tracking studies would provide better understanding of spatial and temporal movements of individuals on Reunion Island and potentially identify currently unknown mating sites (Conenna et al., 2019).

## Conclusion

Our study illustrates how understanding mechanisms involved in speciation can be challenging and thus the importance of integrating past evolutionary processes and contemporary gene flows. Here, we demonstrate that fine-scale sampling scheme and multi-marker comparisons at regional and local scales are necessary to achieve a complete picture of the population structure and history of island endemic bats. Such genetic approaches in combination with understanding a range of ecological parameters are also crucial to reduce uncertainty in conservation decision making of vulnerable mammals due of their endemicity status, such as *Mormopterus francoismoutoui*.

## Acknowledgements

We are grateful to personnel of Eco-Med Océan Indien, Biotope, the Direction de l’Exploitation et de l’Entretien des Routes (DEER) of Région Réunion, the Direction des Routes et des Transports (DRT) of Département Réunion, and the Salazie city hall for their help in identifying and accessing bat roosts. We are thankful to David Wilkinson for fruitful discussions and help with analyses. We also thank Yann Gomard and Julien Mélade for previous laboratory work on Mauritius and Malagasy samples. We also thank Guillaume Verchère for his assistance in the field. This research was supported by the French National Research Agency (ANR JCJC SEXIBAT), by the European Regional Development Funds ERDF PO INTERREG V ECOSPIR number RE6875. Samantha Aguillon was supported by a “Contrat Doctoral de l’Université de La Réunion”.

## Conflict of interest

We have no competing interests to declare.

## Data accessibility and benefits-sharing section

Mitochondrial sequences (D-loop) has been deposited in Genbank under ID’s from OR081945 to OR082607 and microsatellites genotypes and metadata are available at Zenodo (https://doi.org/10.5281/zenodo.8069702). D-loop sequences for the Reunion Island dataset are coded with the roost name, the field identification number and the sex (F = female and M = male). Regional sequences are coded with the island (MADA = Madagascar and MAU = Mauritius) and field identification number.

## Author contributions

S.A. and M.D. designed the study. S.A., C.C., A.D., G.L.M., C.L., A.O.H., C.T., L.J., P.T., S.M.G., and M.D. performed sampling of biological material. S.A., C.C., A.D., M.G., and M.D. generated the data. S.A., C.C., A.D., and M.D. analysed the data. S.A. and M.D. led the writing, with comments and final approval from all co-authors.

## Notes

https://doi.org/10.5281/zenodo.8069702

### Competing Interest Statement

The authors have declared no competing interest.

## References

Aguillon, S., Le Minter, G., Lebarbenchon, C., Hoarau, A. O. G., Toty, C., Joffrin, L., Ramanantsalama, R. V., Augros, S., Tortosa, P., Mavingui, P., & Dietrich, M. (2023). A population in perpetual motion: Highly dynamic roosting behavior of a tropical island endemic bat. Ecology and Evolution, 13(2), e9814. https://doi.org/10.1002/ece3.9814

Akaike, H. (1974). A new look at the statistical model identification. IEEE Transactions on Automatic Control, 19(6), 716–723. https://doi.org/10.1109/TAC.1974.1100705

Ancillotto, L., Fichera, G., Pidinchedda, E., Veith, M., Kiefer, A., Mucedda, M., & Russo, D. (2021). Wildfires, heatwaves and human disturbance threaten insular endemic bats. Biodiversity and Conservation, 30(14), 4401–4416.

Andriollo, T., Naciri, Y., & Ruedi, M. (2015). Two mitochondrial barcodes for one biological species: The case of European Kuhl’s Pipistrelles (Chiroptera). PLoS One, 10(8), e0134881. https://doi.org/10.1371/journal.pone.0134881

Augros, S., Denis, B., Crozet, P., Roué, S.G., & Fabulet, P.-Y. (2015). La cohabitation entre l’homme et les microchiroptères à La Réunion : bilan actualisé, retours d’expérience et outils de conservation. Vespère, 5, 371–384.

Belkhir, K., Borsa, P., Chikhi, L., Raufaste, N. & Bonhomme, F. (2004). GENETIX 4.05, Population genetics software for Windows TM pour la génétique des populations. Laboratoire Génome, Populations, Interactions, CNRS UMR 5000, Université de Montpellier II, Montpellier (France).

Bellard, C., Leclerc, C., & Courchamp, F. (2014). Impact of sea level rise on the 10 insular biodiversity hotspots. Global Ecology and Biogeography, 23(2), 203–212. https://doi.org/10.1111/geb.12093

Biollaz, F., Bruyndonckx, N., Beuneux, G., Mucedda, M., Goudet, J., & Christe, P. (2010). Genetic isolation of insular populations of the Maghrebian bat, *Myotis punicus*, in the Mediterranean Basin. Journal of Biogeography, 37(8), 1557–1569. https://doi.org/10.1111/j.1365-2699.2010.02282.x

Bouckaert, R., Vaughan, T. G., Barido-Sottani, J., Duchêne, S., Fourment, M., Gavryushkina, A., Heled, J., Jones, G., Kühnert, D., Maio, N. D., Matschiner, M., Mendes, F. K., Müller, N. F., Ogilvie, H. A., Plessis, L. du, Popinga, A., Rambaut, A., Rasmussen, D., Siveroni, I., Suchard, M. A., Wu, C.-H., Xie, D., Zhang, C., Stadler, T. & Drummond, A. J. (2019). BEAST 2.5: An advanced software platform for Bayesian evolutionary analysis. PLoS Computational Biology, 15(4), e1006650. https://doi.org/10.1371/journal.pcbi.1006650

Cadet, T. (1980). Données récentes sur l’origine, l’âge et la structure géologique de l’île de La Réunion. Académie de l’île de La Réunion, Bulletin, 1969-1978(24), 73–87.

Calderón-Acevedo, C. A., Rodríguez-Durán, A., & Soto-Centeno, J. A. (2021). Effect of land use, habitat suitability, and hurricanes on the population connectivity of an endemic insular bat. Scientific Reports, 11(1), 1–11.

Conenna, I., López-Baucells, A., Rocha, R., Ripperger, S., & Cabeza, M. (2019). Movement seasonality in a desert-dwelling bat revealed by miniature GPS loggers. Movement Ecology, 7(1), 1–10. https://doi.org/10.1186/s40462-019-0170-8

Darriba, D., & Posada, D. (2016). jModelTest 2 Manual v0. 1.10. Parallel Computing, 9, 772.

Dietrich, M., Minter, G. L., Turpin, M., & Tortosa, P. (2019). Development and characterization of a multiplex panel of microsatellite markers for the Reunion free-tailed bat *Mormopterus francoismoutoui*. PeerJ, 7, e8036. https://doi.org/10.7717/peerj.8036

Dietrich, M., Wilkinson, D. A., Benlali, A., Lagadec, E., Ramasindrazana, B., Dellagi, K., & Tortosa, P. (2015). *Leptospira* and paramyxovirus infection dynamics in a bat maternity enlightens pathogen maintenance in wildlife. Environmental Microbiology, 17(11), 4280–4289. https://doi.org/10.1111/1462-2920.12766

Do, C., Waples, R. S., Peel, D., Macbeth, G. M., Tillett, B., & Ovenden, J. (2013). NeEstimator v2: Re-implementation of software for the estimation of contemporary effective population size (Ne) from genetic data. Molecular Ecology Resources, 14, 209–214. https://doi.org/10.1111/1755-0998.12157

Drummond, A. J., Ho, S. Y. W., Phillips, M. J., & Rambaut, A. (2006). Relaxed phylogenetics and dating with confidence. PLoS Biology, 4(5), e88.

Drummond, A. J., Rambaut, A., Shapiro, B., & Pybus, O. G. (2005). Bayesian coalescent inference of past population dynamics from molecular sequences. Molecular Biology and Evolution, 22(5), 1185–1192.

Estoup, A., Jarne, P., & Cornuet, J.-M. (2002). Homoplasy and mutation model at microsatellite loci and their consequences for population genetics analysis. Molecular Ecology, 11(9), 1591–1604. https://doi.org/10.1046/j.1365-294X.2002.01576.x

Evanno, G., Regnaut, S., & Goudet, J. (2005). Detecting the number of clusters of individuals using the software structure: A simulation study. Molecular Ecology, 14(8), 2611– 2620. https://doi.org/10.1111/j.1365-294X.2005.02553.x

Excoffier, L., & Lischer, H. (2015). ARLEQUIN Ver 3.5: An integrated software package for population genetics data analysis. Swiss Institute of Bioinformatics.

Festa, F., Ancillotto, L., Santini, L., Pacifici, M., Rocha, R., Toshkova, N., Amorim, F., Benítez-López, A., Domer, A., Hamidović, D., Kramer-Schadt, S., Mathews, F., Radchuk, V., Rebelo, H., Ruczynski, I., Solem, E., Tsoar, A., Russo, D., & Razgour, O. (2023). Bat responses to climate change: A systematic review. Biological Reviews, 98(1), 19–33. https://doi.org/10.1111/brv.12893

Frankham, R. (1996). Relationship of genetic variation to population size in wildlife. Conservation Biology, 10(6), 1500–1508.

Fu, Y. X. (1997). Statistical tests of neutrality of mutations against population growth, hitchhiking and background selection. Genetics, 147(2), 915–925.

Gauffre, B., Estoup, A., Bretagnolle, V., & Cosson, J. F. (2008). Spatial genetic structure of a small rodent in a heterogeneous landscape. Molecular Ecology, 17(21), 4619–4629. https://doi.org/10.1111/j.1365-294X.2008.03950.x

Glass, B. P. (1982). Seasonal movements of Mexican freetail bats *Tadarida brasiliensis mexicana* banded in the Great Plains. The Southwestern Naturalist, 27(2), 127–133.

Gomard, Y., Dietrich, M., Wieseke, N., Ramasindrazana, B., Lagadec, E., Goodman, S. M., Dellagi, K., & Tortosa, P. (2016). Malagasy bats shelter a considerable genetic diversity of pathogenic *Leptospira* suggesting notable host-specificity patterns. FEMS Microbiology Ecology, 92(4), fiw037.

Goodman, S. M., van Vuuren, B. J., Ratrimomanarivo, F., Probst, J.-M., & Bowie, R. C. K. (2008). Specific status of populations in the Mascarene Islands referred to *Mormopterus acetabulosus* (Chiroptera: Molossidae), with description of a new species. Journal of Mammalogy, 89(5), 1316–1327. https://doi.org/10.1644/07-MAMM-A-232.1

Goudet, J. (2003). FSTAT (version 2.9. 4), a program (for Windows 95 and above) to estimate and test population genetics parameters. Department of Ecology & Evolution, Lausanne University, Switzerland, 53.

Halczok, T. K., Brändel, S. D., Flores, V., Puechmaille, S. J., Tschapka, M., Page, R. A., & Kerth, G. (2018). Male-biased dispersal and the potential impact of human-induced habitat modifications on the Neotropical bat *Trachops cirrhosus*. Ecology and Evolution, 8(12), 6065–6080. https://doi.org/10.1002/ece3.4161

Harpending, H. C. (1994). Signature of ancient population growth in a low-resolution mitochondrial DNA mismatch distribution. Human Biology, 66(4),591–600.

Harrison, R. G. (1989). Animal mitochondrial DNA as a genetic marker in population and evolutionary biology. Trends in Ecology & Evolution, 4(1), 6–11. https://doi.org/10.1016/0169-5347(89)90006-2

Hasegawa, M., Kishino, H., & Yano, T. (1985). Dating of the human-ape splitting by a molecular clock of mitochondrial DNA. Journal of Molecular Evolution, 22(2), 160–174. https://doi.org/10.1007/BF02101694

Ho, S. Y. W., & Shapiro, B. (2011). Skyline-plot methods for estimating demographic history from nucleotide sequences. Molecular Ecology Resources, 11(3), 423–434. https://doi.org/10.1111/j.1755-0998.2011.02988.x

Hogner, S., Laskemoen, T., Lifjeld, J. T., Porkert, J., Kleven, O., Albayrak, T., Kabasakal, B., & Johnsen, A. (2012). Deep sympatric mitochondrial divergence without reproductive isolation in the common redstart *Phoenicurus phoenicurus*. Ecology and Evolution, 2(12), 2974–2988. https://doi.org/10.1002/ece3.398

Hubisz, M. J., Falush, D., Stephens, M., & Pritchard, J. K. (2009). Inferring weak population structure with the assistance of sample group information. Molecular Ecology Resources, 9(5), 1322–1332.

Jang, J. E., Byeon, S. Y., Kim, H., Kim, J., Myeong, H.-H., & Lee, H. J. (2021). Genetic evidence for sex-biased dispersal and cryptic diversity in the greater horseshoe bat, *Rhinolophus ferrumequinum*. Biodiversity and Conservation, 30, 847–864. https://doi.org/10.1007/s10531-021-02120-y

Joffrin, L., Goodman, S. M., Wilkinson, D. A., Ramasindrazana, B., Lagadec, E., Gomard, Y., Le Minter, G., Dos Santos, A., Corrie Schoeman, M., Sookhareea, R., Tortosa, P., Julienne, S., Gudo, E. S., Mavingui, P., & Lebarbenchon, C. (2020). Bat coronavirus phylogeography in the Western Indian Ocean. Scientific Reports, 10(1), 6873. https://doi.org/10.1038/s41598-020-63799-7

Johnson, L. N. L., McLeod, B. A., Burns, L. E., Arseneault, K., Frasier, T. R., & Broders, H. G. (2015). Population genetic structure within and among seasonal site types in the little brown bat (*Myotis lucifugus*) and the northern long-eared bat (*M. septentrionalis*). PLoS One, 10(5), e0126309. https://doi.org/10.1371/journal.pone.0126309

Jones, K. E., Barlow, K. E., Vaughan, N., Rodríguez-Durán, A., & Gannon, M. R. (2001). Short-term impacts of extreme environmental disturbance on the bats of Puerto Rico. Animal Conservation, 4(1), 59–66. https://doi.org/10.1017/S1367943001001068

Jones, K., Mickleburgh, S., Sechrest, W., & Walsh, A. (2009). Global overview of the conservation of island bats: Importance challenges and opportunities. In T. H. Fleming & P. A. Racey (Eds.), Island bats: Evolution, ecology, and conservation (pp. 496–533). The University of Chicago Press.

Jung, K., & Threlfall, C. G. (2018). Trait-dependent tolerance of bats to urbanization: A global meta-analysis. Proceedings of the Royal Society B: Biological Sciences, 285(1885), 20181222. https://doi.org/10.1098/rspb.2018.1222

Kuo, H.-C., Chen, S.-F., Fang, Y.-P., Cotton, J. A., Parker, J. D., Csorba, G., Lim, B. K., Eger, J. L., Chen, C.-H., Chou, C.-H., & Rossiter, S. J. (2015). Speciation processes in putative island endemic sister bat species: False impressions from mitochondrial DNA and microsatellite data. Molecular Ecology, 24(23), 5910–5926. https://doi.org/10.1111/mec.13425

Lagabrielle, E., Rouget, M., Payet, K., Wistebaar Mahlangu, P., Durieux, L., Baret, S., Lombard, A., & Strasberg, D. (2009). Identifying and mapping biodiversity processes for conservation planning in islands: A case study in Réunion Island (Western Indian Ocean). Biological Conservation, 142, 1523–1535. https://doi.org/10.1016/j.biocon.2009.02.022

Laine, V. N., Sävilammi, T., Wahlberg, N., Meramo, K., Ossa, G., Johnson, J. S., Blomberg, A. S., Yeszhanov, A. B., Yung, V., Paterson, S., & Lilley, T. M. (2023). Whole-genome analysis reveals contrasting relationships among nuclear and mitochondrial genomes between three sympatric bat species. Genome Biology and Evolution, 15(1), evac175. https://doi.org/10.1093/gbe/evac175

Lloyd, B. D. (2003). The demographic history of the New Zealand short-tailed bat *Mystacina tuberculata* inferred from modified control region sequences. Molecular Ecology, 12(7), 1895–1911. https://doi.org/10.1046/j.1365-294X.2003.01879.x

Loureiro, L. O., Engstrom, M. D., & Lim, B. K. (2020). Comparative phylogeography of mainland and insular species of Neotropical molossid bats (*Molossus*). Ecology and Evolution, 10(1), 389–409. https://doi.org/10.1002/ece3.5903

Makhov, I. A., Gorodilova, Y. Y., & Lukhtanov, V. A. (2021). Sympatric occurrence of deeply diverged mitochondrial DNA lineages in Siberian geometrid moths (Lepidoptera: Geometridae): Cryptic speciation, mitochondrial introgression, secondary admixture or effect of *Wolbachia*? Biological Journal of the Linnean Society, 134(2), 342–365.

McCracken, G. F., Gillam, E. H., Westbrook, J. K., Lee, Y.-F., Jensen, M. L., & Balsley, B. B. (2008). Brazilian free-tailed bats (*Tadarida brasiliensis*: Molossidae, Chiroptera) at high altitude: Links to migratory insect populations. Integrative and Comparative Biology, 48(1), 107–118.

McCracken, G. F., & Wilkinson, G. S. (2000). Bat mating systems. In E. G. Crichton & P. H. Krutzsch (Eds.), Reproductive biology of bats (pp. 321–362). Academic Press. https://doi.org/10.1016/B978-012195670-7/50009-6

Mélade, J., Wieseke, N., Ramasindrazana, B., Flores, O., Lagadec, E., Gomard, Y., Goodman, S. M., Dellagi, K., & Pascalis, H. (2016). An eco-epidemiological study of Morbilli-related paramyxovirus infection in Madagascar bats reveals host-switching as the dominant macro-evolutionary mechanism. Scientific Reports, 6(1), 1–12.

Moussy, C., Hosken, D., Aegerter, J., Mathews, F., Smith, G., & Bearhop, S. (2013). Migration and dispersal patterns of bats and their influence on genetic structure. Mammal Review, 43, 183–195. https://doi.org/10.1111/j.1365-2907.2012.00218.x

Naidoo, T., Schoeman, M. C., Goodman, S. M., Taylor, P. J., & Lamb, J. (2016). Discordance between mitochondrial and nuclear genetic structure in the bat *Chaerephon pumilus* (Chiroptera: Molossidae) from southern Africa. Mammalian Biology - Zeitschrift Fur Saugetierkunde, 81(2), 115–122. https://doi.org/10.1016/j.mambio.2015.11.002

Nehlig, P., & Marie, B. (2005). Connaissance géologique de la Réunion - Livret de l’enseignant. BRGM Editions;

Oyler-McCance, S., Fike, J., Lukacs, P., Sparks, D., O’Shea, T., & Whitaker Jr, J. O. (2017). Genetic mark–recapture improves estimates of maternity colony size for Indiana bats. Journal of Fish and Wildlife Management, 9(1), 25–35. https://doi.org/10.3996/122016-JFWM-093

Peterson, R. L. (1985). Systematic review of the molossid bats allied with the genus *Mormopterus* (Chiroptera: Molossidae). Acta Zoologica Fennica, 170, 205–208.

Petit, E., Excoffier, L., & Mayer, F. (1999). No evidence of bottleneck in the postglacial recolonization of Europe by the Noctule Bat (*Nyctalus noctula*). Evolution, 53(4), 1247–1258. https://doi.org/10.1111/j.1558-5646.1999.tb04537.x

Pinzari, C. A., Bellinger, M. R., Price, D., & Bonaccorso, F. J. (2023). Genetic diversity, structure, and effective population size of an endangered, endemic hoary bat, ʻōpeʻapeʻa, across the Hawaiian Islands. PeerJ, 11, e14365. https://doi.org/10.7717/peerj.14365

Pritchard, J. K., Wen, X., & Falush, D. (2010). Documentation for structure software: Version 2.3. University of Chicago, Chicago, IL, 1–37.

Rambaut, A., & Drummond, A. J. (2015). LogCombiner v1. 8.2. LogCombinerv1, 8, 656. Rambaut, A., & Drummond, A. J. (2019). TreeAnnotator v 2 6.0-MCMC output analysis. Software Development. Part of Beast, 2.

Rambaut, A., Drummond, A. J., Xie, D., Baele, G., & Suchard, M. A. (2018). Posterior summarization in Bayesian phylogenetics using Tracer 1.7. Systematic Biology, 67(5), 901–904.

Ratrimomanarivo, F. H., Goodman, S. M., Taylor, P. J., Melson, B., & Lamb, J. (2009). Morphological and genetic variation in *Mormopterus jugularis* (Chiroptera: Molossidae) in different bioclimatic regions of Madagascar with natural history notes. Mammalia, 73(2), 110–129. https://doi.org/10.1515/MAMM.2009.032

Reardon, T. B., McKenzie, N. L., Cooper, S. J. B., Appleton, B., Carthew, S., Adams, M. (2014). A molecular and morphological investigation of species boundaries and phylogenetic relationships in Australian free-tailed bats *Mormopterus* (Chiroptera: Molossidae). Australian Journal of Zoology, 62, 109–136.

Richardson, J. L., Michaelides, S., Combs, M., Djan, M., Bisch, L., Barrett, K., Silveira, G., Butler, J., Aye, T. T., Munshi-South, J., DiMatteo, M., Brown, C., & McGreevy Jr, T. J. (2021). Dispersal ability predicts spatial genetic structure in native mammals persisting across an urbanization gradient. Evolutionary Applications, 14(1), 163–177. https://doi.org/10.1111/eva.13133

Rivers, N. M., Butlin, R. K., & Altringham, J. D. (2006). Autumn swarming behaviour of Natterer’s bats in the UK: Population size, catchment area and dispersal. Biological Conservation, 127(2), 215–226. https://doi.org/10.1016/j.biocon.2005.08.010

Rodhouse, T. J., Rodriguez, R. M., Banner, K. M., Ormsbee, P. C., Barnett, J., & Irvine, K. M. (2019). Evidence of region-wide bat population decline from long-term monitoring and Bayesian occupancy models with empirically informed priors. Ecology and Evolution, 9(19), 11078–11088. https://doi.org/10.1002/ece3.5612

Rogers, A. R., & Harpending, H. (1992). Population growth makes waves in the distribution of pairwise genetic differences. Molecular Biology and Evolution, 9(3), 552–569.

Rozas, J., Ferrer-Mata, A., Sánchez-DelBarrio, J. C., Guirao-Rico, S., Librado, P., Ramos-Onsins, S. E., & Sánchez-Gracia, A. (2017). DnaSP 6: DNA sequence polymorphism analysis of large data sets. Molecular Biology and Evolution, 34(12), 3299–3302. https://doi.org/10.1093/molbev/msx248

Rstudio Team. (2021). Integrated development for R. RStudio, Inc. https://www.rstudio.com/products/rstudio

Russo, D., & Ancillotto, L. (2015). Sensitivity of bats to urbanization: A review. Mammalian Biology, 80(3), 205–212.

Salinas-Ramos, V. B., Ancillotto, L., Bosso, L., Sánchez-Cordero, V., & Russo, D. (2020). Interspecific competition in bats: State of knowledge and research challenges. Mammal Review, 50(1), 68–81. https://doi.org/10.1111/mam.12180

Sandron, F. (2007). Dynamique de la population réunionnaise (1663-2030). In F. Sandron (Eds.), La population réunionnaise. Analyse démographique (pp. 27–41). IRD Editions.

Silva, S., Ferreira, G., Pamplona, H., Carvalho, T., Cordeiro, J., & Trevelin, L. (2020). Effects of landscape heterogeneity on population genetic structure and demography of Amazonian phyllostomid bats. Mammal Research, 66, 217–225. https://doi.org/10.1007/s13364-020-00546-3

Smouse, R. P. P., & Peakall, R. (2012). GenAlEx 6.5: Genetic analysis in Excel. Population genetic software for teaching and research—an update. Bioinformatics, 28(19), 2537–2539.

Sun, K., Kimball, R. T., Liu, T., Wei, X., Jin, L., Jiang, T., Lin, A., & Feng, J. (2016). The complex evolutionary history of big-eared horseshoe bats (*Rhinolophus macrotis* complex): Insights from genetic, morphological and acoustic data. Scientific Reports, 6(1), 35417. https://doi.org/10.1038/srep35417

Taki, Y., Vincenot, C. E., Sato, Y., & Inoue-Murayama, M. (2021). Genetic diversity and population structure in the Ryukyu flying fox inferred from remote sampling in the Yaeyama archipelago. PLoS ONE, 16(3), e0248672. https://doi.org/10.1371/journal.pone.0248672

Tamura, K., & Nei, M. (1993). Estimation of the number of nucleotide substitutions in the control region of mitochondrial DNA in humans and chimpanzees. Molecular Biology and Evolution, 10(3), 512–526.

Toews, D. P. L., & Brelsford, A. (2012). The biogeography of mitochondrial and nuclear discordance in animals. Molecular Ecology, 21(16), 3907–3930. https://doi.org/10.1111/j.1365-294X.2012.05664.x

Van Oosterhout, C., Hutchinson, W. F., Wills, D. P., & Shipley, P. (2004). MICRO-CHECKER: Software for identifying and correcting genotyping errors in microsatellite data. Molecular Ecology Notes, 4(3), 535–538.

Warren, B. H., Simberloff, D., Ricklefs, R. E., Aguilée, R., Condamine, F. L., Gravel, D., Morlon, H., Mouquet, N., Rosindell, J., Casquet, J., Conti, E., Cornuault, J., Fernández-Palacios, J. M., Hengl, T., Norder, S. J., Rijsdijk, K. F., Sanmartín, I., Strasberg, D., Triantis, K. A., Valente, L. M., Whittaker, R. J., Gillespie, R. G., Emerson, B. C., & Thébaud, C. (2015). Islands as model systems in ecology and evolution: Prospects fifty years after MacArthur-Wilson. Ecology Letters, 18(2), 200–217. https://doi.org/10.1111/ele.12398

Webb, W. C., Marzluff, J. M., & Omland, K. E. (2011). Random interbreeding between cryptic lineages of the common raven: Evidence for speciation in reverse. Molecular Ecology, 20(11), 2390–2402. https://doi.org/10.1111/j.1365-294X.2011.05095.x

Weir, B. S., & Cockerham, C. C. (1984). Estimating F-statistics for the analysis of population structure. Evolution, 38(6), 1358–1370.

Yule, G. U. (1925). A mathematical theory of evolution, based on the conclusions of Dr. J. C. Willis, F.R.S. *Philosophical Transactions of the Royal Society of London. Series B*, Containing Papers of a Biological Character, 213, 21–87.

