## Supplementary Material for "Stuck on a small tropical island: wide *in-situ* diversification of an urban-dwelling bat"

**Table S1 Summary of mitochondrial and microsatellite data used in this study for *Mormopterus francoismoutoui*.**

Maternity roosts are indicated with an asterisk (*) next to the roost name. The first number indicates the total number of sequences, with females and males in parentheses and the sites are indicated in Figure 1.

| Roost | Habitat | Pregnancy 2018 (summer) | Non-reproductive 2019 (winter) | Mating 2019 – 2020 (March) | Opportunistic sampling | | Total |
|---|---|---|---|---|---|---|---|
| | | | | | February 2019 | October 2018 | |
| **AOM** | Building | | | 14 (8/6) | | | 14 |
| | | | | 30 (15/15) | | | 30 |
| **CIT** | Building | 15 (0/15) | | | | | 15 |
| | | 29 (0/29) | | | | | 29 |
| **EGI** | Building | 26 (8/18) | | 15 (8/7) | | | 41 |
| | | 34 (11/23) | | 30 (15/15) | | | 64 |
| **ESA** | Bridge | 15 (9/6) | 16 (9/7) | 15 (8/7) | | | 46 |
| | | 36 (19/16) | 34 (18/16) | 26 (19/7) | | | 96 |
| **MON** | Bridge | 18 (3/15) | 15 (7/8) | 15 (8/7) | | | 48 |
| | | 33 (3/30) | 30 (15/15) | 25 (16/9) | | | 88 |
| **PBV** | Cliff | | | | 11 (4/7) | | 11 |
| | | | | | 11 (4/7) | | 11 |
| **PSR** | Building | 17 (6/11) | 15 (2/13) | 15 (7/8) | | | 47 |
| | | 34 (12/22) | 34 (2/32) | 28 (7/21) | | | 96 |
| **RAC** | Bridge | | | | | 16 (4/12) | 16 |
| | | | | | | 14 (4/10) | 14 |
| **RBL** | Bridge | 16 (8/8) | 15 (7/8) | 15 (7/8) | | | 46 |
| | | 31 (13/18) | 34 (11/23) | 28 (20/8) | | | 93 |
| **RES** | Bridge | 15 (7/8) | | 15 (7/8) | | | 30 |
| | | 32 (15/17) | | 30 (7/23) | | | 62 |
| **RPQ** | Bridge | 15 (6/9) | 15 (8/7) | 15 (8/7) | | | 45 |
| | | 37 (19/18) | 33 (11/22) | 30 (15/15) | | | 100 |
| **STJ** | Building | | 15 (8/7) | | | | 15 |

| | | | | | | | |
|---|---|---|---|---|---|---|---|
| | | | 30 (13/17) | | | | 30 |
| **STM** | Bridge | 15 (7/8) | 13 (3/10) | 15 (3/12) | | | 43 |
| | | 34 (17/17) | 13 (3/10) | 23 (3/20) | | | 70 |
| **TBA*** | Cave | 31 (29/2) | | 15 (7/8) | | | 46 |
| | | 34 (31/3) | | 29 (22/7) | | | 63 |
| **TGI*** | Building | 19 (12/7) | 15 (7/8) | 15 (7/8) | | | 49 |
| | | 38 (20/18) | 36 (13/23) | 29 (15/14) | | | 103 |
| **TM5** | Bridge | | | | 15 (7/8) | | 15 |
| | | | | | 28 (13/15) | | 28 |
| **TRI** | Bridge | 15 (1/14) | 15 (8/7) | | | | 30 |
| | | 29 (1/28) | 29 (9/20) | | | | 58 |
| **VSP** | Bridge | 16 (7/9) | 15 (7/8) | 15 (7/8) | | | 46 |
| | | 38 (18/20) | 35 (7/28) | 28 (15/13) | | | 101 |
| **Total** | | 233 | 149 | 179 | 26 | 16 | 603 |
| | | 439 | 308 | 336 | 39 | 14 | 1136 |

**Table S2 Genetic diversity indices calculated with mitochondrial DNA (D-loop) of *Mormopterus francoismoutoui* separately for females and males.**

N is the number of sequences used for calculation, Hd is the haplotype diversity, and π is the nucleotide diversity. Roost sites are defined in Table S1.

| Roost | Females | | | Males | | |
|---|---|---|---|---|---|---|
| | N | Hd | π | N | Hd | π |
| **AOM** | 8 | 0.964 | 0.0256 | 6 | 1.000 | 0.0288 |
| **CIT** | / | / | / | 15 | 1.000 | 0.0309 |
| **EGI** | 16 | 1.000 | 0.0310 | 25 | 0.997 | 0.0274 |
| **ESA** | 26 | 1.000 | 0.0263 | 20 | 0.995 | 0.0284 |
| **MON** | 18 | 1.000 | 0.0304 | 30 | 1.000 | 0.0278 |
| **PBV** | 4 | 1.000 | 0.0251 | 7 | 1.000 | 0.0336 |
| **PSR** | 15 | 1.000 | 0.0293 | 33 | 0.998 | 0.0307 |
| **RAC** | 4 | 1.000 | 0.0249 | 12 | 1.000 | 0.0298 |
| **RBL** | 22 | 1.000 | 0.0292 | 24 | 1.000 | 0.0280 |
| **RES** | 14 | 1.000 | 0.0290 | 16 | 1.000 | 0.0292 |
| **RPQ** | 22 | 1.000 | 0.0297 | 23 | 0.992 | 0.0298 |
| **STJ** | 8 | 1.000 | 0.0258 | 7 | 1.000 | 0.0298 |
| **STM** | 13 | 1.000 | 0.0295 | 30 | 1.000 | 0.0284 |
| **TBA** | 36 | 0.998 | 0.0292 | 10 | 1.000 | 0.0303 |
| **TGI** | 26 | 0.997 | 0.0313 | 23 | 0.996 | 0.0292 |
| **TM5** | 7 | 1.000 | 0.0307 | 8 | 1.000 | 0.0305 |
| **TRI** | 9 | 1.000 | 0.0296 | 21 | 1.000 | 0.0284 |
| **VSP** | 21 | 1.000 | 0.0286 | 25 | 0.997 | 0.0299 |
| **TOTAL** | 269 | 0.998 | 0.0290 | 334 | 0.998 | 0.0282 |

**Table S3 Genetic diversity indices calculated with mitochondrial DNA (D-loop) of *Mormopterus francoismoutoui* captured during the pregnancy, non-reproductive, and supposed mating periods.**

N is the number of sequences used for calculation, Hd is the haplotype diversity, and π is the nucleotide diversity. Roost sites are defined in Table S1.

| Roost | Pregnancy 2018 (summer) | | | Non-reproductive 2019 (winter) | | | Mating 2019 – 2020 (March) | | |
|---|---|---|---|---|---|---|---|---|---|
| | N | Hd | π | N | Hd | π | N | Hd | π |
| **AOM** | / | / | / | / | / | / | 14 | 0.989 | 0.0264 |
| **CIT** | 15 | 1.000 | 0.0296 | / | / | / | / | / | / |
| **EGI** | 26 | 1.000 | 0.0288 | / | / | / | 15 | 0.990 | 0.0289 |
| **ESA** | 15 | 0.990 | 0.0264 | 16 | 1.000 | 0.0277 | 15 | 0.990 | 0.0279 |
| **MON** | 18 | 1.000 | 0.0291 | 15 | 1.000 | 0.0296 | 15 | 1.000 | 0.0293 |
| **PBV** | / | / | / | / | / | / | / | / | / |
| **PSR** | 17 | 1.000 | 0.0304 | 15 | 0.990 | 0.0306 | 15 | 1.000 | 0.0306 |
| **RAC** | / | / | / | / | / | / | / | / | / |
| **RBL** | 16 | 0.992 | 0.0305 | 15 | 1.000 | 0.0293 | 15 | 1.000 | 0.0279 |
| **RES** | 15 | 1.000 | 0.0270 | / | / | / | 15 | 1.000 | 0.0306 |
| **RPQ** | 15 | 0.990 | 0.0295 | 15 | 1.000 | 0.0298 | 15 | 1.000 | 0.0313 |
| **STJ** | / | / | / | 15 | 1.000 | 0.0278 | / | / | / |
| **STM** | 15 | 1.000 | 0.0317 | 13 | 1.000 | 0.0287 | 15 | 1.000 | 0.0280 |
| **TBA** | 31 | 0.998 | 0.0299 | / | / | / | 15 | 1.000 | 0.0272 |
| **TGI** | 19 | 1.000 | 0.0329 | 15 | 1.000 | 0.0294 | 15 | 1.000 | 0.0292 |
| **TM5** | / | / | / | / | / | / | / | / | / |
| **TRI** | 15 | 1.000 | 0.0279 | 15 | 1.000 | 0.0310 | / | / | / |
| **VSP** | 16 | 1.000 | 0.0286 | 15 | 1.000 | 0.0268 | 15 | 1.000 | 0.0304 |
| **TOTAL** | 233 | 0.998 | 0.0287 | 149 | 0.998 | 0.0287 | 179 | 0.998 | 0.0285 |

**Table S4 Genetic diversity indices calculated with 11 microsatellite markers of *Mormopterus francoismoutoui* separately for females and males.**

N is the number of genotypes used, Ho is the observed heterozygosity, and He is the expected heterozygosity. Standard error is given following ±. The roost sites are defined in Table S1.

| Roost | Females | | | Males | | |
|---|---|---|---|---|---|---|
| | N | Ho | He | N | Ho | He |
| **AOM** | 15 | 0.781 ± 0.039 | 0.776 ± 0.033 | 15 | 0.743 ± 0.038 | 0.776 ± 0.038 |
| **CIT** | / | / | / | 29 | 0.787 ± 0.035 | 0.791 ± 0.029 |
| **EGI** | 26 | 0.750 ± 0.047 | 0.793 ± 0.033 | 38 | 0.777 ± 0.039 | 0.807 ± 0.027 |
| **ESA** | 57 | 0.767 ± 0.033 | 0.802 ± 0.033 | 39 | 0.788 ± 0.038 | 0.803 ± 0.033 |
| **MON** | 34 | 0.795 ± 0.040 | 0.785 ± 0.033 | 54 | 0.797 ± 0.030 | 0.801 ± 0.036 |
| **PBV** | 4 | 0.803 ± 0.062 | 0.721 ± 0.036 | 7 | 0.732 ± 0.045 | 0.738 ± 0.025 |
| **PSR** | 21 | 0.799 ± 0.034 | 0.785 ± 0.032 | 75 | 0.784 ± 0.039 | 0.795 ± 0.037 |
| **RAC** | 4 | 0.894 ± 0.064 | 0.688 ± 0.043 | 10 | 0.814 ± 0.077 | 0.773 ± 0.033 |
| **RBL** | 44 | 0.768 ± 0.026 | 0.793 ± 0.038 | 49 | 0.758 ± 0.032 | 0.802 ± 0.032 |
| **RES** | 22 | 0.743 ± 0.052 | 0.801 ± 0.030 | 40 | 0.744 ± 0.036 | 0.793 ± 0.035 |
| **RPQ** | 45 | 0.795 ± 0.039 | 0.799 ±0.034 | 55 | 0.782 ± 0.030 | 0.803 ± 0.033 |
| **STJ** | 13 | 0.800 ± 0.027 | 0.766 ± 0.034 | 17 | 0.769 ± 0.037 | 0.793 ± 0.036 |
| **STM** | 23 | 0.829 ± 0.042 | 0.792 ± 0.036 | 47 | 0.782 ± 0.042 | 0.808 ± 0.031 |
| **TBA** | 53 | 0.788 ± 0.038 | 0.799 ± 0.031 | 10 | 0.756 ± 0.062 | 0.746 ± 0.057 |
| **TGI** | 48 | 0.765 ± 0.049 | 0.804 ± 0.034 | 55 | 0.769 ± 0.045 | 0.802 ± 0.035 |
| **TM5** | 12 | 0.727 ± 0.051 | 0.769 ± 0.033 | 16 | 0.772 ± 0.048 | 0.777 ± 0.034 |
| **TRI** | 10 | 0.742 ± 0.054 | 0.757 ± 0.029 | 48 | 0.790 ± 0.037 | 0.803 ± 0.031 |
| **VSP** | 40 | 0.789 ± 0.036 | 0.800 ± 0.036 | 61 | 0.773 ± 0.030 | 0.800 ± 0.033 |
| **TOTAL** | 471 | 0.784 ± 0.011 | 0.778 ± 0.008 | 665 | 0.773 ± 0.010 | 0.789 ± 0.008 |

**Table S5 Genetic diversity indices calculated with 11 microsatellite markers of *Mormopterus francoismoutoui* captured during pregnancy/summer, non-reproductive/winter period, and supposed mating period.**

N is the number of genotypes used, Ho is the observed heterozygosity, and He is the expected heterozygosity. Standard error is given following ±. Roost sites are defined in Table S1.

| Roost | Pregnancy 2018 (summer) | | | Non-reproductive 2019 (winter) | | | Mating 2019 – 2020 (March) | | |
|---|---|---|---|---|---|---|---|---|---|
| | N | Ho | He | N | Ho | He | N | Ho | He |
| AOM | / | / | / | / | / | / | 30 | 0.762 ± 0.035 | 0.793 ± 0.037 |
| CIT | 29 | 0.787 ± 0.035 | 0.791 ± 0.029 | / | / | / | / | / | / |
| EGI | 34 | 0.783 ± 0.029 | 0.808 ± 0.025 | / | / | / | 30 | 0.748 ± 0.063 | 0.794 ± 0.036 |
| ESA | 36 | 0.798 ± 0.025 | 0.809 ± 0.034 | 34 | 0.757 ± 0.048 | 0.788 ± 0.032 | 26 | 0.768 ± 0.038 | 0.796 ± 0.032 |
| MON | 33 | 0.796 ± 0.033 | 0.800 ± 0.033 | 30 | 0.804 ± 0.031 | 0.791 ± 0.034 | 25 | 0.784 ± 0.049 | 0.777 ± 0.039 |
| PBV | / | / | / | / | / | / | / | / | / |
| PSR | 34 | 0.796 ± 0.033 | 0.797 ± 0.031 | 34 | 0.781 ± 0.040 | 0.775 ± 0.041 | 28 | 0.784 ± 0.046 | 0.792 ± 0.036 |
| RAC | / | / | / | / | / | / | / | / | / |
| RBL | 31 | 0.764 ±0.038 | 0.783 ± 0.037 | 34 | 0.760 ± 0.033 | 0.797 ± 0.034 | 28 | 0.773 ± 0.027 | 0.800 ± 0.031 |
| RES | 32 | 0.745 ± 0.035 | 0.791 ± 0.035 | / | / | / | 30 | 0.744 ± 0.048 | 0.795 ± 0.032 |
| RPQ | 37 | 0.783 ± 0.039 | 0.795 ± 0.038 | 33 | 0.780 ± 0.036 | 0.788 ± 0.034 | 30 | 0.802 ± 0.038 | 0.804 ± 0.030 |
| STJ | / | / | / | 30 | 0.782 ± 0.029 | 0.796 ± 0.034 | / | / | / |
| STM | 34 | 0.794 ± 0.039 | 0.803 ± 0.032 | 13 | 0.846 ± 0.051 | 0.794 ± 0.033 | 23 | 0.774 ± 0.050 | 0.794 ± 0.031 |
| TBA | 34 | 0.799 ± 0.039 | 0.791 ± 0.033 | / | / | / | 29 | 0.764 ± 0.048 | 0.797 ± 0.035 |
| TGI | 38 | 0.783 ± 0.040 | 0.792 ± 0.037 | 36 | 0.762 ± 0.057 | 0.798 ± 0.030 | 29 | 0.748 ± 0.052 | 0.801 ± 0.040 |
| TM5 | / | / | / | / | / | / | / | / | / |
| TRI | 29 | 0.772 ± 0.035 | 0.803 ± 0.030 | 29 | 0.789 ± 0.048 | 0.784 ± 0.035 | / | / | / |
| VSP | 38 | 0.798 ± 0.035 | 0.790 ± 0.035 | 35 | 0.770 ± 0.036 | 0.799 ± 0.035 | 28 | 0.766 ± 0.046 | 0.797 ± 0.031 |
| TOTAL | 439 | 0.784 ± 0.009 | 0.796 ± 0.009 | 308 | 0.783 ± 0.013 | 0.791 ± 0.010 | 336 | 0.768 ± 0.013 | 0.795 ± 0.009 |

**Table S6 Pairwise differentiation between roost among global population of *Mormopterus francoismoutoui*** with mitochondrial DNA values (Φst) below the diagonal and microsatellite markers (Fst) above. No significant p-values after Holm correction. Roost sites are defined in Table S1.

| Roost | AOM | CIT | EGI | ESA | MON | PBV | PSR | RAC | RBL | RES | RPQ | STM | STJ | TBA | TGI | TM5 | TRI | VSP |
|---|---|---|---|---|---|---|---|---|---|---|---|---|---|---|---|---|---|---|
| **AOM** | - | 0.018 | 0.011 | -0.017 | 0.017 | 0.006 | 0.026 | 0.022 | -0.0005 | 0.015 | 0.015 | 0.003 | 0.043 | 0.033 | 0.006 | -0.018 | 0.0008 | 0.011 |
| **CIT** | -0.002 | - | -0.001 | 0.009 | -0.006 | -0.027 | 0.015 | -0.020 | -0.002 | 0.002 | 0.001 | -0.002 | 0.006 | 0.0007 | -0.016 | -0.013 | -0.003 | -0.003 |
| **EGI** | -0.0008 | -0.003 | - | -0.0006 | -0.010 | -0.031 | 0.0016 | -0.025 | -0.003 | -0.0005 | -0.013 | -0.015 | 0.010 | -0.0002 | -0.004 | -0.020 | -0.009 | -0.004 |
| **ESA** | 0.002 | 0.004 | -0.0009 | - | 0.004 | 0.0008 | 0.023 | -0.004 | -0.0007 | -0.003 | 0.007 | -0.004 | 0.018 | 0.011 | 0.010 | -0.025 | -0.0007 | -0.0007 |
| **MON** | 0.0006 | 0.001 | -0.00005 | 0.002 | - | -0.030 | 0.010 | -0.029 | -0.008 | 0.001 | -0.008 | -0.008 | 0.011 | -0.001 | -0.005 | -0.014 | -0.008 | -0.010 |
| **PBV** | 0.003 | -0.007 | 0.0007 | 0.006 | 0.004 | - | -0.026 | -0.040 | -0.020 | -0.007 | -0.035 | -0.025 | -0.012 | -0.022 | -0.028 | -0.033 | -0.032 | -0.023 |
| **PSR** | -0.00002 | -0.0006 | -0.001 | 0.003 | 0.0002 | 0.003 | - | -0.013 | 0.0129 | 0.004 | -0.005 | 0.003 | -0.004 | -0.0001 | 0.004 | 0.005 | 0.010 | 0.005 |
| **RAC** | 0.008 | -0.007 | 0.0009 | 0.005 | 0.005 | -0.003 | 0.003 | - | -0.018 | -0.016 | -0.022 | -0.021 | -0.008 | -0.019 | -0.018 | -0.023 | -0.021 | -0.022 |
| **RBL** | -0.0008 | -0.0006 | -0.0009 | 0.002 | 0.0001 | 0.004 | 0.002 | 0.002 | - | -0.007 | -0.003 | -0.006 | 0.006 | -0.001 | -0.009 | -0.016 | -0.008 | -0.013 |
| **RES** | -0.0008 | -0.0009 | -0.001 | -0.0007 | 0.0009 | 0.003 | -0.0002 | -0.0003 | -0.0003 | - | -0.008 | -0.003 | -0.017 | -0.016 | -0.003 | -0.012 | 0.005 | -0.011 |
| **RPQ** | -0.002 | -0.002 | -0.0009 | 0.0006 | 0.001 | 0.003 | 0.0008 | 0.004 | 0.00003 | -0.0002 | - | -0.012 | -0.003 | -0.010 | -0.005 | -0.015 | -0.007 | -0.009 |
| **STM** | -0.001 | -0.004 | -0.001 | 0.002 | -0.00003 | 0.003 | 0.001 | -0.0006 | -0.001 | -0.0009 | -0.001 | - | 0.005 | -0.001 | -0.006 | -0.022 | -0.011 | -0.008 |
| **STJ** | 0.002 | -0.005 | -0.0006 | 0.005 | 0.004 | -0.001 | 0.0007 | 0.0006 | 0.002 | -0.002 | 0 | -0.002 | - | -0.018 | -0.0004 | -0.006 | 0.008 | -0.006 |
| **TBA** | 0.0005 | -0.002 | -0.003 | -0.001 | -0.002 | 0.005 | -0.002 | -0.002 | -0.001 | -0.004 | -0.001 | -0.0008 | 0.0004 | - | 0.0002 | -0.010 | 0.0004 | -0.011 |
| **TGI** | 0.002 | -0.003 | -0.0006 | 0.001 | 0.0001 | 0.0002 | 0.0004 | 0.003 | 0.001 | 0.0002 | -0.0007 | -0.001 | 0.001 | -0.002 | - | -0.015 | -0.0007 | -0.006 |
| **TM5** | 0.004 | 0.005 | 0.003 | 0.005 | 0.003 | 0.017 | 0.003 | 0.010 | 0.003 | 0.003 | 0.003 | 0.005 | 0.008 | 0.006 | 0.004 | - | -0.025 | -0.015 |
| **TRI** | -0.001 | -0.001 | -0.002 | 0.001 | -0.002 | 0.005 | -0.001 | 0.004 | -0.001 | -0.002 | -0.0003 | -0.0002 | 0.001 | -0.003 | -0.001 | 0.002 | - | -0.007 |
| **VSP** | 0.0007 | -0.0002 | -0.001 | 0.0008 | -0.0005 | 0.008 | -0.001 | 0.003 | -0.0003 | -0.002 | -0.0004 | -0.0004 | 0.0002 | -0.002 | -0.0003 | 0.003 | -0.001 | - |

**Table S7 Summary of isolation by distance tests for each sex and season for both mtDNA and microsatellite markers in *Mormopterus francoismoutoui*.** r is the regression coefficient and *p* is the significance of the test.

| | **Mitochondrial data** | | **Microsatellites data** | |
|---|---|---|---|---|
| | r | *p* | r | *p* |
| **Females** | 0.06 | 0.27 | -0.03 | 0.63 |
| **Males** | -0.12 | 0.88 | 0.009 | 0.44 |
| **Pregnancy 2018 (summer)** | -0.03 | 0.536 | 0.03 | 0.42 |
| **Non-reproductive 2019 (winter)** | 0.21 | 0.12 | 0.06 | 0.39 |
| **Mating 2019 – 2020 (March)** | -0.26 | 0.97 | -0.15 | 0.87 |

**Table S8 Summary of effective sample size (ESS) for each dimension group parameters of *Mormopterus francoismoutoui* implemented in the Coalescent Bayesian Skyline model from mitochondrial data (D-loop).**

The model with five groups (in red) have the best likelihood ESS followed by six and ten group models.

| Dimension group | Posterior ESS | Likelihood ESS | Prior ESS | Bayesian Skyline ESS |
|---|---|---|---|---|
| **3** | 2290 | 1160 | 3497 | 3499 |
| **4** | 2501 | 816 | 3726 | 3576 |
| **5** | 3073 | **1634** | 3354 | 3200 |
| **6** | 2480 | 1618 | 3069 | 2929 |
| **7** | 3030 | 1010 | 3623 | 3400 |
| **8** | 2382 | 1130 | 3336 | 2789 |
| **9** | 2543 | 1168 | 4035 | 3717 |
| **10** | 2745 | 1617 | 3253 | 3042 |

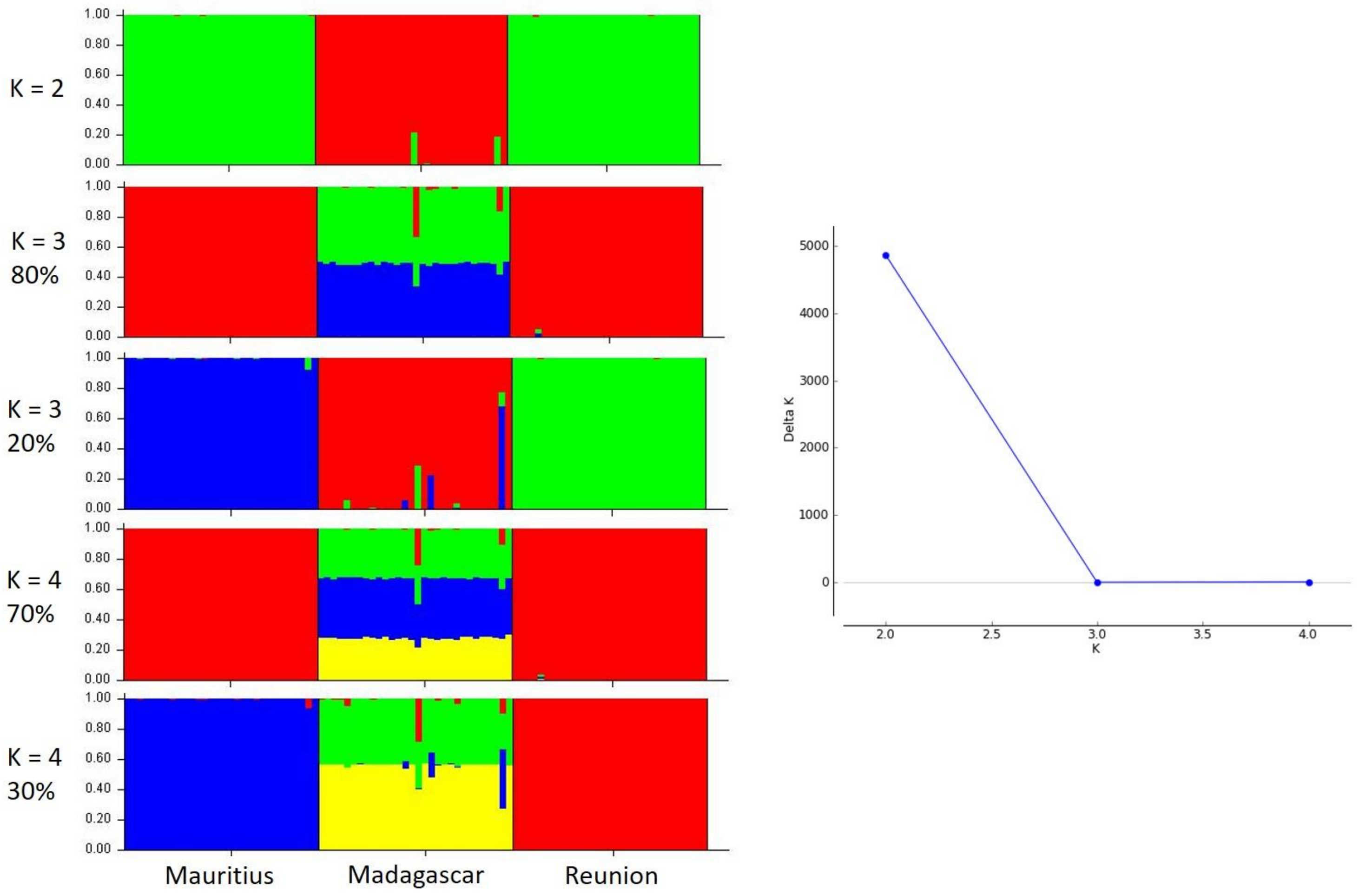


**Figure S1 Assignment plots among regional *Mormopterus* performed in STRUCTURE for K from 2 to 4.**

The best K (K = 2) was determined following the Evanno method (Delta K). Each vertical bar represents one individual and colours indicates genetic clusters. Islands are indicated below the plot and percentages runs are indicated below the K.

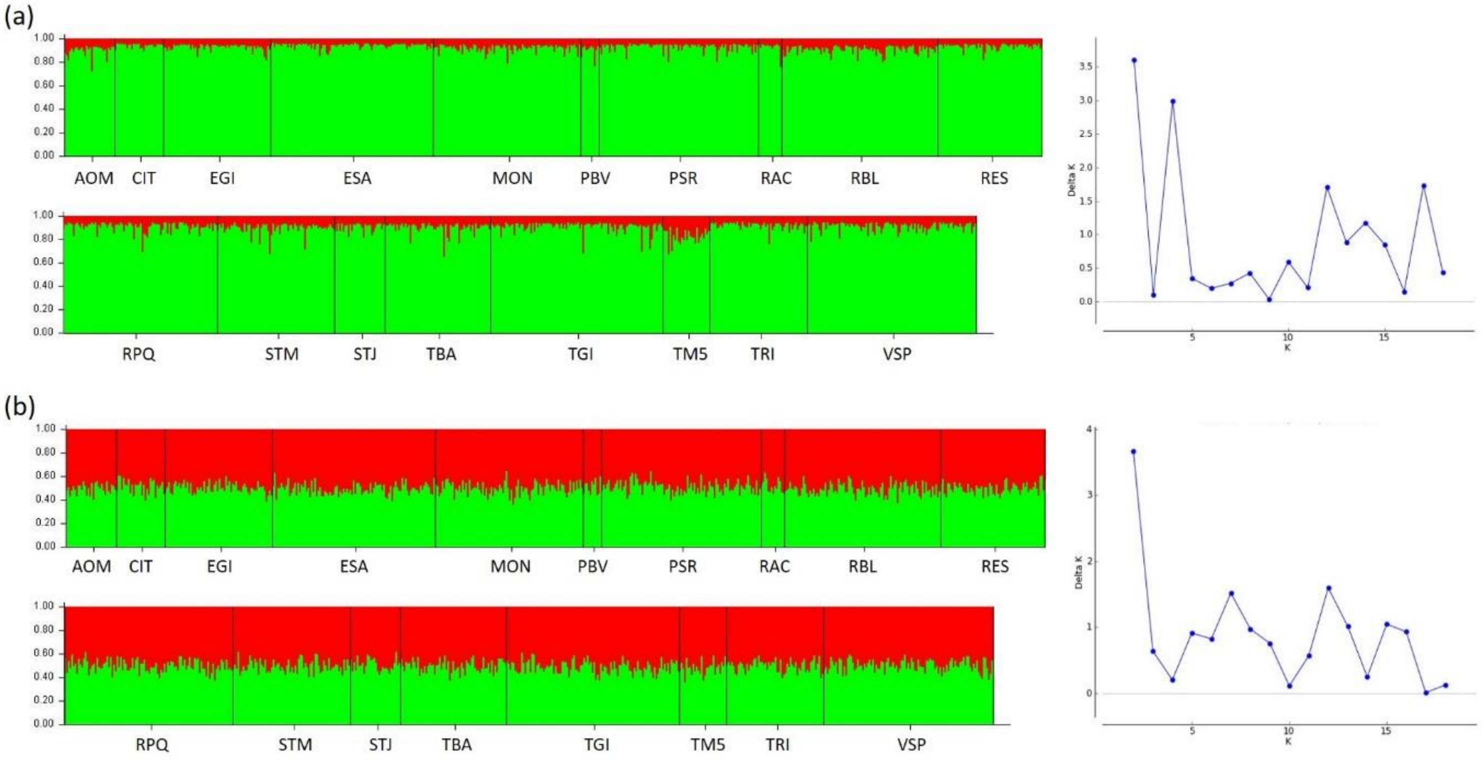


**Figure S2 Assignment plot among Reunion population of *Mormopterus francoismoutoui* in STRUCTURE for the best K =2 following the Evanno Method (Delta K).**

Results are shown (a) with Locprior and (b) without Locprior model.

Each vertical bar represents one individual and colours indicate genetic clusters. Roost sites are indicated below the plot and these are defined in Table S1.

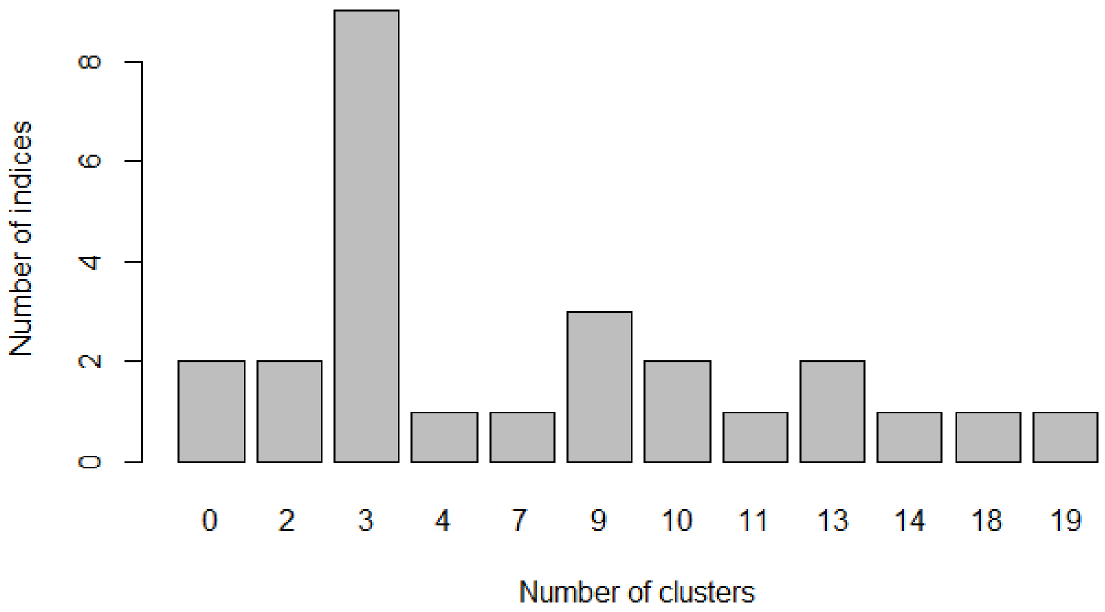


**Figure S3 Bar plot showing the best number of genetic clusters of *Mormopterus francoismoutoui* according to 26 indices evaluated with the k-means clustering.**

The greatest frequency of indices found the best number of clusters being K = 3.
